# Sex-biased *Yap1* oncogene function

**DOI:** 10.1101/2022.12.21.521494

**Authors:** Nourhan Abdelfattah, Sivaraman Natarajan, Jose Maldonado, Han Nhat Tran, Rachael McMinimy, Hannah Borland, Shu-hsia Chen, Fernando Camargo, James Olson, Joshy George, Kyuson Yun

## Abstract

The incidence of many human cancers differs according to sex, but little is known about the interplay between oncogenic events and sex as a variable in tumorigenesis. Here we report that the oncogene *Yap1* is sexually dimorphic in medulloblastoma progression and immune suppression. We show that *Yap1* promotes stemness and blocks differentiation in sonic hedgehog (SHH)-subtype medulloblastoma by at least two distinct but complementary molecular mechanisms to regulate the RNA expression and protein functions of *Sox2, Atoh1, NeuroD1, and Zic1/2. Yap1* also promotes an immune suppressive tumor microenvironment by directly regulating *Csf1, Igf1,* and *Igfbp3* transcription and modulating IL6-JAK-STAT3, TNFR1, TGF-β, and CCL5 immune pathways. Notably, *Yap1* function is more critical in males and this is evolutionarily conserved: genes downstream of YAP1 identified in mouse models stratify male but not female medulloblastoma patient survival. In summary, we demonstrate a sex-based function for an oncogene, underscoring the critical need to incorporate sex as a variable in cancer mechanism and clinical response studies, particularly those involving YAP1.

## INTRODUCTION

There is a known sex bias in cancer incidence and survival^1^, and recent studies have shown that this bias also extends to therapy responses, including immunotherapies^2–4^. However, the underlying mechanisms are poorly understood. Although sex hormones play obvious and significant roles in the pathogenesis of some tumors such as estrogen receptor-positive breast or androgen receptor-positive prostate cancers, they do not explain the clear sex bias seen in many other cancers, such as brain cancers^5–7^. Cell-autonomous mechanisms such as epigenetic changes during organogenesis and aging, and X-chromosome-linked gene expression contribute to sexual dimorphism^8–10^. There are also reports of sexually dimorphic roles for important tumor suppressor genes such as *TP53* and *RB* in tumorigenesis^10, 11^. However, little is known about the sex-specific roles played by oncogenes in tumorigenesis.

Male sex is a well-recognized, but understudied, risk factor for medulloblastoma (MB), and MB incidence is consistently higher in males than females across all subgroups and ages^12–16^. Moreover, for patients older than three years of age, females have significantly better outcomes than males^17, 18^. MB is a primitive neuroectodermal tumor and the most common pediatric brain malignancy, comprising ∼20% of primary pediatric tumors and accounting for ∼10% of pediatric cancer-related deaths^19, 20^. Major developments in clinical care, including risk stratification, molecular subtyping, and aggressive multimodal treatments, have considerably increased five-year overall survival (OS) rates for MB patients^19, 20^. Nevertheless, ∼20-30% of tumors are incurable^19^, and aggressive treatments can lead to severe long-term sequelae including neurocognitive deficits, neuropathy, and secondary malignancies^19, 21^. Immunotherapies – now considered the fourth pillar of cancer care – can provide durable therapeutic benefits while minimizing devastating side-effects but, despite >55 FDA-approved immunotherapies for other cancers, there is no FDA-approved immunotherapy for MB^22, 23^. This is in part because MB is considered an immune “cold” tumor with an immune suppressive microenvironment containing very few T cells^22, 24^. There is therefore considerable interest in developing strategies to promote T cell infiltration into MB and other immune-cold tumors.

MB is stratified into four distinct molecular subtypes: WNT, sonic hedgehog (SHH), group 3, and group 4^20, 25, 26^. Human SHH MBs are identified by activating mutations in or high expression of SHH pathway genes such as *PTCH* or *SMO*^20^. Genomic analyses of human MB have revealed that Yes-associated protein 1 (*YAP1)* is amplified or over-expressed in a subset of SHH MBs^27^. The Hippo/Yap pathway is one of the top ten most commonly dysregulated pathways in human cancers^28, 29^, and it is an essential signaling pathway for stem cell homeostasis and organ size control^30^. YAP1 is a downstream transcriptional effector of the Hippo pathway that regulates downstream target genes by binding to co-factors such as TEAD^31, 32^ to promote the expression of genes that control stemness, tumorigenesis, and invasion/metastasis in mouse models and human cancers^28, 33–38^. More recent studies have shown that *YAP1* also plays an important but complicated role in tumor immunity^39^. For example, a cell-autonomous function for YAP1 in Tregs is necessary for the immune suppressive Treg phenotype^35^. YAP1 has been also shown to activate PD-L1 expression in thoracic cancer^40^ and BRAF inhibitor-resistant melanomas, dampening CD8^-^ T cell-mediated immune responses^34^. In prostate^37^ and pancreatic cancer^41^, YAP1 has been shown to recruit myeloid-derived suppressor cells (MDSCs) to promote tumor progression.

Here we investigated the functional significance of *Yap1* amplification and overexpression in SHH-MB *in vivo*. We discovered an unexpected male bias in *Yap1* function in MB formation in mice. Mechanistically, YAP1 directly represses expression of transcription factors that promote the differentiation of cerebellar neural precursor cells (*NeuroD1, Zic1/2)* and activates expression of *Sox2*, which promotes normal neural and MB stem cells^42^. In addition, YAP1 promotes tumor immune evasion by regulating transcription of cytokines and chemokines such as *Csf1, Igf1,* and *Igfbp3* that modulate immune infiltrates in MB. Surprisingly YAP1 binding regions in mouse MB cells *in vivo* are not enriched for TEAD binding sites, suggesting a non-canonical transcriptional regulation mechanism. Finally, we corroborate an evolutionarily conserved male bias for YAP1 and its downstream target genes in human MBs.

## RESULTS

### Yap1 is reactivated in SHH MBs regardless of cell-of-origin

Since *Yap1* is preferentially amplified or overexpressed in a subset of SHH MBs, we first tested whether *Yap1* expression in SHH MB depends on the developmental stage of different cells-of-origin for MB transformation. To do so, we generated four different SHH MB mouse models by activating the SHH pathway in neural stem cells (NSCs) and neural progenitors at different stages of maturation (**Supplementary Fig. 1a,b**). *Ptch1^+/−^* mice harbor a germline mutation in the *Ptch1* gene and develop spontaneous medulloblastomas arising from NSCs or cerebellar granule neuron progenitor cells (cGNPs), with or without cooperating *Trp53* mutations^43, 44^. *fSMO-M2* mice harbor a floxed allele of the constitutively active *Smoothened* (*SMO-M2*) transgene that activates the SHH pathway upon Cre-mediated recombination^45^. When crossed to *Olig2-cre* mice, the transgene is expressed in NSCs and oligodendrocyte precursors and motor neurons^46^. 100% of *fSMO-M2;Olig2-cre* mice develop MBs with average survival of 57 days^46, 47^ . When crossed to hGFAP-cre mice, *fSMO-M2;hGFAP-cre* mice (SG) express the *SMO-M2* transgene in embryonic NSCs and their progeny. 100% of SG mice develop MBs by weaning age^44, 47^. *NeuroD2-SmoA1* mice express the constitutively active point mutation transgene *SmoA1* under the control of the neurogenic differentiation 2 (*Neurod2*) promoter in maturing cGNPs, and 48% of these mice develop spontaneous MB^48^ (**Supplementary Fig. 1a-c)**.

Immunohistochemical analysis with a YAP1 antibody in each of these SHH MB models showed YAP1 expression in SHH-induced MB cells in all four models (**Supplementary Fig. 1c-d**). YAP1 expression in SHH MB is therefore independent of cell-of-origin. Next, we analyzed YAP1 expression in patient-derived xenograft (PDX) models of MB. YAP1 was robustly expressed in SHH PDX model MED1712 in both the cytoplasm and nucleus but not in group 3 PDX model MED211 (**Supplementary Fig. 1e**), consistent with previous reports identifying YAP1 protein staining as a histological marker of human SHH MB^49^. Our metanalysis of three published human MB bulk RNA-seq datasets revealed higher *YAP1* expression in both WNT and SHH MBs compared with group 3 and group 4 MBs (**Supplementary Fig. 1f**). Furthermore, our analysis of a publicly available human MB single-cell RNA-sequencing dataset^50^ (GSE119926) (**Supplementary Fig. 1g,h, Supplementary Table 1**) showed that 5-20% of MB cells from patients with adult and infantile SHH MBs as well as WNT subtype MBs express *YAP1* (**Supplementary Fig. 1i**). Together, these data indicate that YAP1 activation is an evolutionarily conserved mechanism in mouse and human SHH MBs.

### Yap1 is required for SMO-induced MB progression but not initiation

To directly test whether *Yap1* is a cooperating oncogene necessary for SHH*-*induced MB, we crossed *Yap1*^f/f^ mice to the *fSMO-M2;hGFAP-Cre* (SG) spontaneous MB mouse model (**Supplementary Fig. 2a**) to selectively and concurrently delete *Yap1* in *SMO-M2*-expressing cells. In *fYap1;fSMO-M2;hGFAPcre* (YSG) mice, *Yap1* is deleted in the same cells that express the *SMO-M2* transgene. In our hands, 100% of SG mice died by 31 days, with a median survival of 22 days (**Fig. 1a**) due to severe hydrocephalus and MB, as previously reported^44^. In contrast, 23 of 40 (57.5%) YSG mice developed considerably milder hydrocephalus and survived significantly longer (35 days to 6 months, p<0.0001, **Fig. 1a and Supplementary Fig. 2b**). These *in vivo* genetic models indicate that *Yap1* is an important cooperating oncogene for SHH-induced MB.

**Figure 1:**
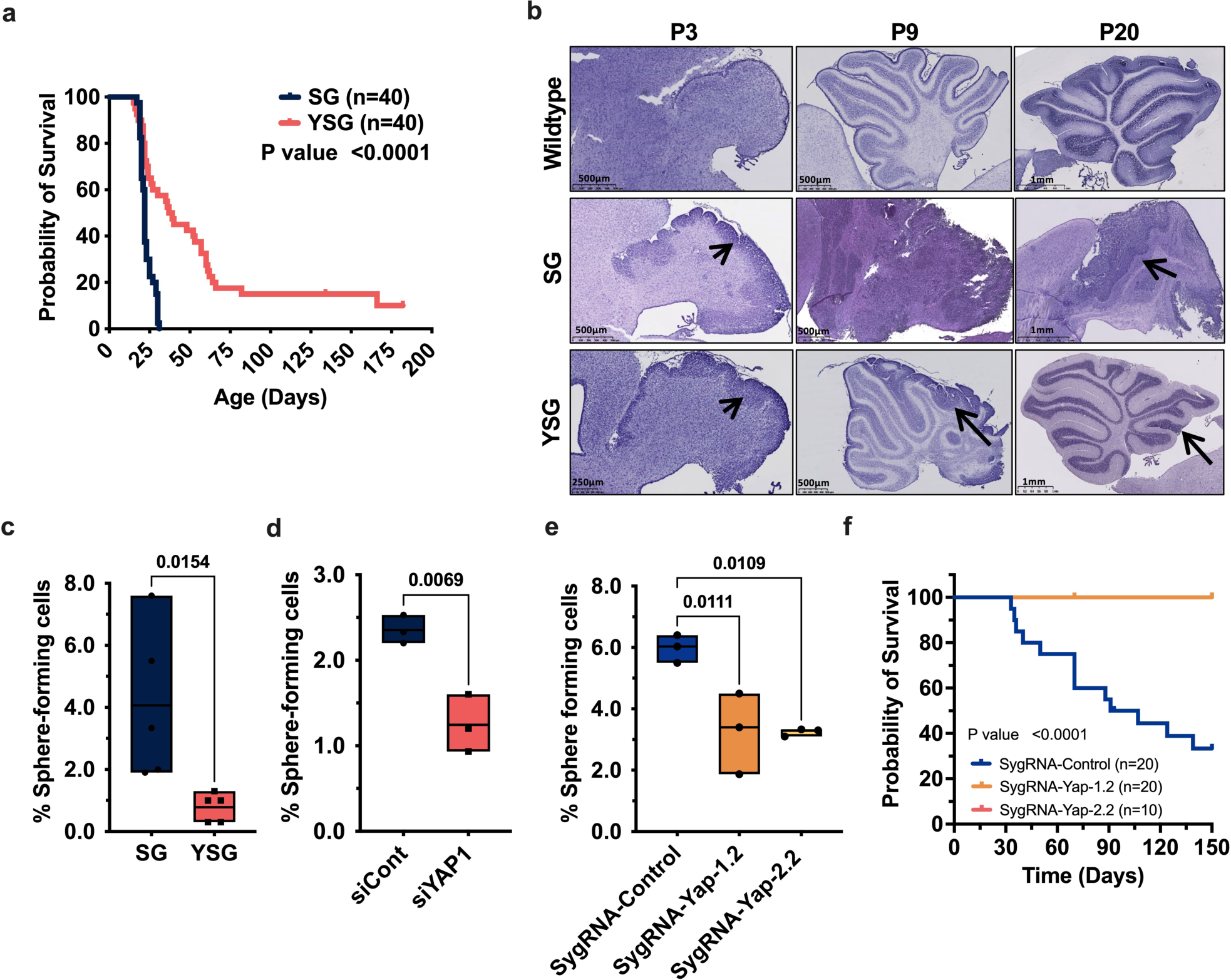
*Yap1* is required for SMO-induced MB progression but not initiation. **a.** Kaplan-Meier survival curves of SG and YSG mice. *P*-value was calculated using the log-rank (Mantel-Cox) test. **b.** H&E staining of wildtype, *SmoM2;hGFAP-cre*(SG), and rescued *Yap1(f/f);SmoM2; hGFAP-cre*(YSGr) brains at p3, p9, and p20. Arrows point to hyperplastic regions. **c.** Secondary sphere-forming assay with tumor spheres isolated from SG and YSG MB. n=5 each. *P*-values were calculated using a two-tailed Student’s t-test. **d.** Self-renewal assay with tumor spheres isolated from primary *Ptch;p53* tumor line treated with Control siRNA or *Yap1* siRNA. n=3 each. *P*-values(e-f) were calculated using ratio paired t-test. **e.** Self renewal assay with tumor spheres isolated from *Ptch;p53* CRISPR-Cas9 clones, control sygRNA vs two different sygRNAs targeting *Yap1*. n=3 each. *P*-values were calculated using one-way ANOVA followed by Dunnett’s multiple comparisons test. **f**. Kaplan-Meier survival curves of B6 mice intracranially injected with control and Yap1 KO *Ptch;p53* cells. *P*-value was calculated using the log-rank (Mantel-Cox) test.

To determine whether *Yap1* is required for *SMO-M2* mediated MB initiation, we analyzed postnatal day 3 (p3), postnatal day 9 (p9), and postnatal day 20 (p20) SG and YSG cerebella. H&E analyses of p3 pups showed equivalent hyperplasia induced by *SMO-M2* expression in SG and YSG pups (arrowheads in **Fig. 1b**), indicating that *Yap1* is not necessary for SMO-induced hyperproliferation and tumor initiation. In contrast, analyses of older animals at p9 and p20 showed normalization of cerebellar development in ∼58% of YSG pups, as visualized by histologically distinguishable cerebellar folia (**Fig. 1b**). Immunohistochemical analysis revealed a reduced density of proliferating Ki67^+^ cells in YSG cerebella (Supplementary Fig. 2c). However, residual hyperplastic EGL regions were present in these rescued YSG (YSG^r^) animals (arrows in **Fig. 1b, lower panel**), eventually giving rise to MBs in these mice. Henceforth, we refer to YSG mice surviving longer than 31 days as YSG^r^ (rescued) mice, and those living less than 31 days as YSG^nr^ (non-rescued) to distinguish these two groups of YSG mice. These results suggests that *Yap1* is dispensable for MB initiation but promotes MB progression (**Fig. 1b**).

To evaluate whether *Yap1* is required for fully transformed MB cell maintenance in an independent model, we isolated and knocked out *Yap1* in primary tumorspheres from established *Ptch1;p53* MBs using a CRISPR-Cas9 system and two different guide RNAs (**Supplementary Fig. 2d**). *Yap1*-KO significantly reduced tumorsphere viability compared with control sygRNA (**Supplementary Fig. 2e**). Additionally, cell cycle analysis of early passage *Yap1*-KO clones showed a significant increase in the frequency of G1 arrested cells compared with control sygRNA (**Supplementary Fig. 2f**).

Considering its known function in stem cell homeostasis in multiple organ systems, we hypothesized that *Yap1* may be necessary to maintain long-term self-renewing cancer stem cells (CSCs), or tumor-initiating cells, in SHH MB. To test this hypothesis, we isolated and measured the self-renewal ability of SG and YSG MB tumorspheres *in vitro.* Primary tumorspheres were isolated from SG and YSG^r^ brains, and YSG cells showed a significant reduction in secondary sphere-forming ability compared with SG tumorspheres (**Fig. 1c**). Similarly, *Yap1* depletion in *Ptch;p53* MBs by either *Yap1* siRNA treatment or CRISPR/Cas9-mediated knockout significantly reduced secondary sphere-forming cells in *Ptch;p53* MB cells (**Fig. 1d,e**), suggesting that *Yap1* function is necessary to maintain MB CSCs. To further test this hypothesis, we orthotopically injected B6 mice with *Ptch;p53* control- or *Yap1*-KO cells to measure tumor initiation *in vivo*. Consistent with the *in vitro* results, none of the mice injected with *Yap1*-KO clones developed MB (*P*-value= 0.0002, **Fig. 1f**), confirming that *Yap1* is required to maintain SHH MB CSCs *in vitro* and *in vivo*.

### YAP1 regulates inflammation, stem cell pathways, and cerebellar neurogenesis

To gain mechanistic insights into how *Yap1* promotes MB CSC maintenance and MB progression, we performed bulk RNA-seq analysis to compare expression profiles of SG and YSG^r^ MBs (**Fig. 2 and Supplementary Table 2**). 1876 genes were significantly differentially expressed between SG and YSG samples (FDR <0.05, FC >2, **Fig. 2a,b**). We performed gene set visualization analysis (GSVA) (**Fig. 2c**), gene ontology (GO) annotation (**Fig. 2d**) and ingenuity pathway analysis (IPA) with differentially expressed genes (**Supplementary Fig. 3a**). As anticipated, Hippo signaling, Notch signaling, and WNT *β*-catenin signaling, critical pathways for NSC maintenance, were among the top 15 downregulated GO biological process annotations (**Fig. 2d**) and GSVA Hallmark dataset analysis (**Fig. 2c**), consistent with loss of stemness in *Yap1*-deleted MBs (**Fig. 2d**). These results suggest that *Yap1* promotes self-renewal and maintenance of MB stem cells through WNT and Notch pathway activation^51, 52^. Unexpectedly, YSG^r^ tumors also showed enrichment of proinflammatory response signatures (interferon gamma response, TNF*α* signaling, complement, inflammatory response, and allograft rejection), activation of cytokine signaling, and leukocyte recruitment and activation pathways (**Fig. 2c,d; Supplementary Fig. 3a**). Upstream regulator analysis using IPA predicted activation of proinflammatory upstream regulators (IFN-γ, TNF, IL1B, IL17, and LPS) and inhibition of immune suppressive regulators such as IL10RA (**Fig. 2e**), suggesting that YSG^r^ MBs are significantly more inflamed than SG MBs.

**Figure 2:**
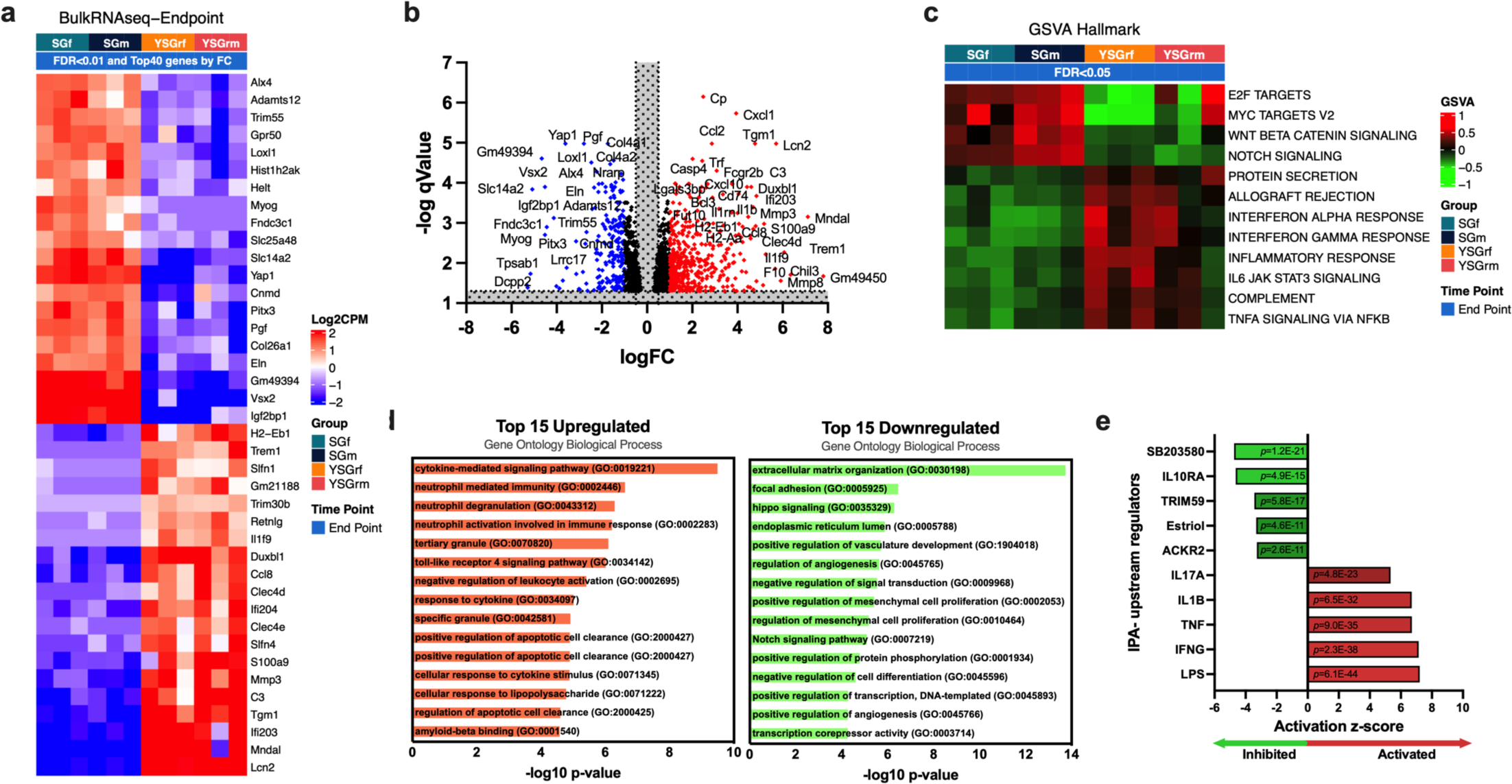
YAP1 regulates inflammation, stem cell pathways, and cerebellar neurogenesis. **a.** Heatmap showing row-centered log-CPM of the top 20 up and down regulated genes between YSG^r^ and SG tumors at experiment endpoint. **b.** Volcano plot highlighting the most significant up or down-regulated genes in endpoint YSG^r^ vs SG MBs. **c**. Gene Set Visualization Analysis (GSVA) of differentially enriched Hallmark pathways between endpoint YSG^r^ and SG tumors (FDR<0.05). **d**. The top 15 up-regulated or downregulated Gene Ontology (GO) biological process annotations sorted by statistical significance. **e**. Ingenuity Pathway Analysis (IPA) prediction of upstream regulators in all YSG^r^ vs SG samples (shown are top and bottom 5 regulators). *P*–value of overlap is written inside the bars.

To determine which of these changes are early, primary effects of *Yap1* deletion in MB cells, we performed bulk RNA-seq analysis of SG, YSG^nr^, and YSG^r^ cerebella from p8 pups (the timepoint at which we can first observe increased neuronal differentiation in YSG^r^ but not in YSG^nr^ and SG brains) (**Supplementary Table 3**). YSG^r^ samples showed similar reductions in cell proliferation hallmarks and WNT *β*-catenin signaling and increased TNF*α* and IL6-JAK-STAT3 signaling gene sets at p8. These results indicate that the molecular changes observed in end-stage YSG^r^ samples, including changes in inflammatory response, start early (**Supplementary Fig. 3b**), and *Yap1* suppresses an inflammatory response in early MBs.

### Yap1 expression in MB cells promotes immune evasion

Considering the strong proinflammatory response molecular signatures in YSG MBs (**Fig. 2**), we hypothesized that *Yap1* expression in MB cells may directly suppress immune cell infiltration. To test this hypothesis, we analyzed the immune phenotypes of SG and YSG MBs by flow cytometry. SG MBs had a significant increase in CD45^hi^ immune infiltrates (i.e., peripheral blood immune cells) in rescued (YSG^r^) mice compared with SG mice or non-rescued YSG^nr^ MB (**Fig. 3a-g, Supplementary Fig. 4**). This included bone marrow-derived monocytes (BMDM: CD45^high^, CD11b^high^, P2RY12^-^) and lymphocytes (CD45^high^, CD11b^-^) (**Supplementary Fig. 4a-b**). To further resolve the different immune subsets, we analyzed the expression of 15+ cell surface immune markers using flow cytometry (**see Methods and Supplementary Fig. 4a-d**). YSG^r^ tumors contained significantly more CD4^+^ and CD8^+^ tumor-infiltrating lymphocytes (TILs) (**Fig. 3b-d**). In addition, neutrophils (CD45^high^, CD11b^high^, Ly6g^+^, Ly6c^int^, SSc), monocytes (CD45^high^, CD11b^high^, Ly6g^lo^, Ly6c^hi^), and macrophages (CD45^high^, CD11b^high^, CD68^+^, P2RY12^neg^, CD11c^-^, Ly6g^lo^, Ly6c^lo^) were also increased (**Fig. 3d,f,g, Supplementary Fig. 4 a,c**). Conversely, the relative fraction of microglia (CD45^int^, CD11b^+^, P2RY12^+^) among total CD45^+^ cells was reduced in YSG^r^ MBs (**Fig. 3d,f,g, Supplementary Fig. 4 a,c**).

**Figure 3:**
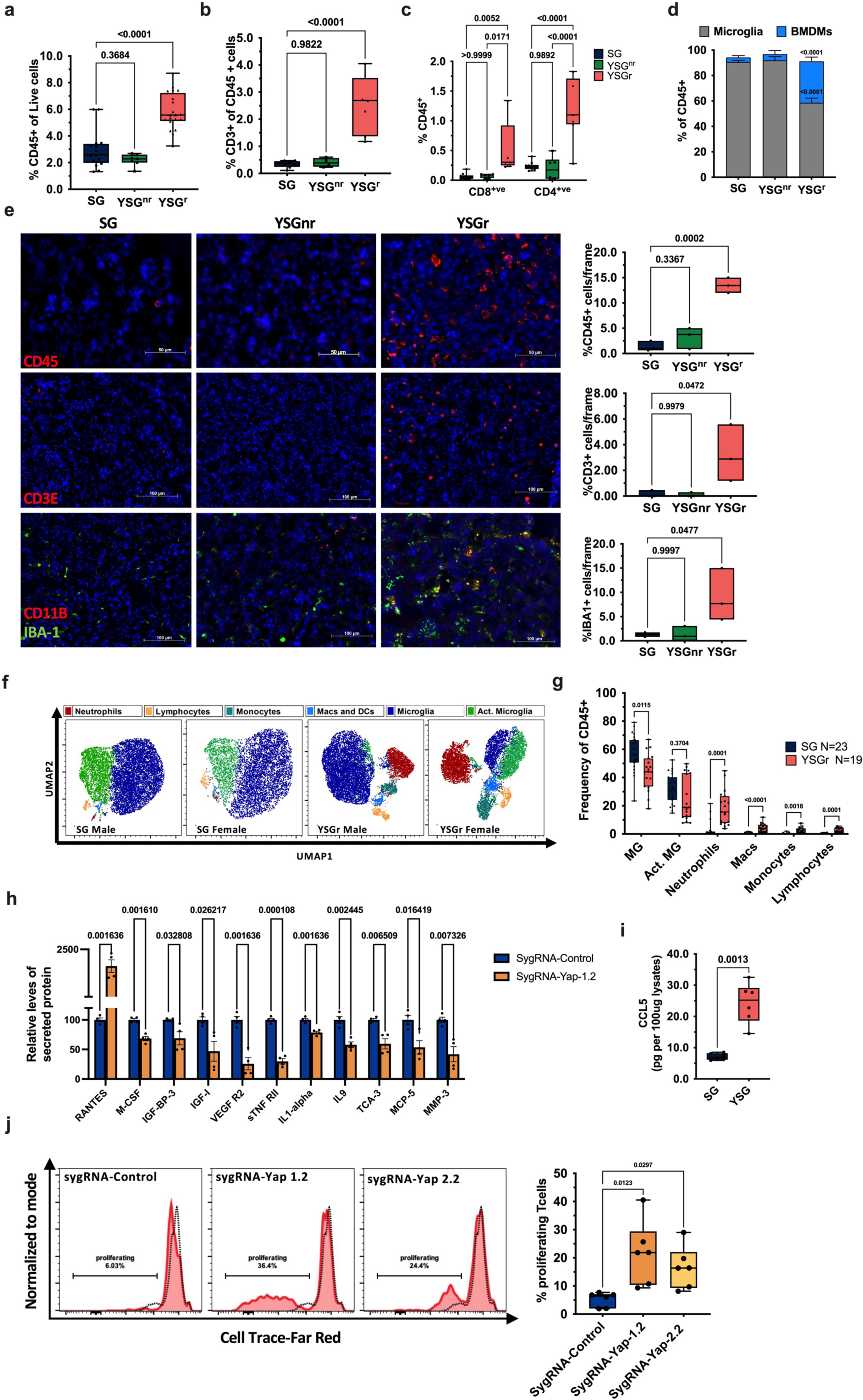
*Yap1* expression in MB cells blocks immune infiltration. **a-d.** Flow cytometry analysis of tumor-infiltrating immune cells shows a significant increase in CD45^+^ cells (SG n=13, YSG^nr^ n=7, YSG^r^ n=13) (a), CD3^+^ cells (SG n=8, YSG^nr^ n=4, YSG^r^ n=7) (b), CD4^+^ and CD8a^+^ cells (SG n=10, YSG^nr^ n=6, YSG^r^ n=7), (c) and BMDMs (SG n=19, YSG^nr^ n=5, YSG^r^ n=14) (d). *P*-values were calculated using one-way ANOVA followed by Dunnett’s multiple comparisons test (a,b) or two-way ANOVA followed by Šídák’s multiple comparisons test (c,d). **e**. Representative immunofluorescence analysis of SG and YSG tumors confirming flow cytometry results. Experiments were performed in triplicate. *P*-values represent Student’s *t*-tests. **f-g.** Flow cytometry analysis of tumor-infiltrating immune subsets. UMAP representation of 10^4^ CD45^+^ cells (f). Quantification of overall tumor-infiltrating immune subsets in SG and YSG mice (g). Line inside box represents median values. *P*-values were calculated using two-way ANOVA followed by Šídák’s multiple comparisons test. See also Supplementary Figure 4. **h.** Secreted cytokine levels were measured in control or Yap KO *Ptch;p53* cell culture supernatants using the RayBio^®^ Mouse Cytokine Antibody Array (n=4).Q-values represent multiple *t*-tests with two-stage step-up (Benjamini, Krieger, and Yekutieli) tests (only significantly secreted cytokines are shown). **i.** Mouse CCL5/RANTES protein levels measured by ELISA (n=6, 3 male and 3 female tumors each). *P*-values were calculated using Student’s *t-*test. **j.** T cell proliferation assay using dye dilution. T cells were isolated from B6 spleens, labeled, and cocultured with control or Yap1 KO *Ptch;p53 cells* for 4 days. Dotted lines represent unstimulated T cells. *P*-value from one-way ANOVA followed by Dunnett’s multiple comparisons test. Whiskers represent minimum and maximum values, the line inside the box represents the mean, and the box extends from the 25th to 75th percentiles. n = 3 (male) and n = 3 (female) B6 spleens.

Since almost all SG mice die by weaning age and weaning alters the immune landscape of developing mice^53^, we tested whether the altered BMDM and T cell infiltration could be due to the older age of rescued YSG mice. We analyzed the immune microenvironment of age-matched SG (from rare SG pups that survived to p27-p31; **Fig 1a**) and YSG MBs (**Supplementary Fig. 5a,b**). In age-matched samples, YSG MB showed significantly higher neutrophil, macrophage, monocyte, and T cell infiltrates compared with SG (**Supplementary Fig. 5a-b**), indicating that *Yap1* deletion-mediated increases in BMDM and T cell infiltration were not a function of age.

An earlier study reported that *Yap1* deletion in CD4^+^ cells reduces T-reg differentiation/function and enhances TIL activation^35^. To rule out the possibility that the increased immune infiltration in YSG^r^ MBs was due to leaky/nonspecific expression of hGFAPcre and *Yap1* deletion in immune cells, we crossed *ROSA26-fEGFP* reporter mice with *hGFAPcre* mice. Flow cytometric analysis of blood, bone marrow, spleen, and cerebella of *ROSA26-fEGFP;hGFAPcre* mice showed no EGFP expression in CD45^+^ immune cells (**Supplementary Fig. 5c**). EGFP expression was restricted to cerebellar CD45^-^ cells (**Supplementary Fig. 5c**), indicating that the observed differences in immune infiltration are not due to *Yap1* deletion in immune cells but rather due to loss of *Yap1* in MB cells.

To assess how *Yap1*-expressing MB cells affect immune cell infiltration, we performed cytokine array analyses of *Ptch;p53* control or *Yap1*-KO tumorsphere culture supernatants to measure the secretion of 96 proteins. Eleven proteins were significantly differentially secreted by *Yap1*-KO tumorsphere cells compared with control-KO cells: M-CSF, IGF-BP-3, IGF-1, VEGF R2, sTNF RII, IL1-*α*, IL9, TCA-3, MCP-5, and MMP-3 were significantly decreased in *Yap1*-KO cell supernatants compared with controls (**Fig. 3h**). In contrast, CCL5/RANTES, a chemotactic agent known to recruit T cells^54^, monocytes^54^, and neutrophils^55^, showed an over 18-fold increase in *Yap1*-KO cell supernatants compared with control-KO cells (**Fig. 3h**). Additionally, CCL5 levels were significantly increased in YSG^r^ tumor lysates compared with SG tumor lysates (**Fig. 3i**). Analysis of our bulk RNA-seq data also showed 57 differentially-expressed chemokines and cytokines between SG and YSG^r^ MBs (FDR <0.05, **Supplementary Fig. 6a**). Together, our evidence shows that *Yap1* directly or indirectly regulates expression of chemokines and cytokines that recruit and polarize T cells, monocytes, and neutrophils in MB.

To test whether *Yap1* expression in MB cells regulates TIL function, we performed a dye dilution assay to measure T cell proliferation/activation in coculture. Freshly isolated B6 splenic naive T cells were cocultured with *Ptch;p53 Yap1*-KO cells or control-KO cells. T cells co-cultured with *Yap1*-KO MB cells showed significantly increased activation/proliferation compared with T cells cocultured with control-KO cells (**Fig 3j**). Thus, changes in the immune landscape in YSG MBs is due to changes in cytokine and chemokine secretion by MB tumor cells, and *Yap1* promotes immune evasion from the very early stages of tumorigenesis.

### Yap1 function is sex biased

During our analysis of immune populations in SG and YSG mice, we observed that there were more steady-state microglia in tumors from female SG mice than male SG mice (*P* = 0.0004, **Fig. 4a**), while male SG tumors had more neutrophil infiltration than female tumors (*P* = 0.0531, **Fig. 4a**). Female YSG MBs showed a significant decrease in the percentage of microglia per CD45^+^ cell compared with female SG MBs (*P* = 0.0004, **Fig. 4b**). In contrast, there was no difference in the percentage of microglia per CD45^+^ cell between male SG and YSG MBs (*P*-value=0.287, **Fig. 4c**). Given the known differences in inflammation and microglia activation states between males and females^56, 57^ and the role of *Yap1* in chemokine/cytokine expression^41^, we examined whether Yap1 exerts a sex-biased function during MB formation. First, we analyzed YSG survival by separating males and females. Intriguingly, 76% of YSG males survived longer than 31 days but only 31% of YSG females showed the rescued phenotype (*P* = 0.015, **Fig. 4d**). Moreover, male YSG mice survived significantly longer than female YSG mice (*P* = 0.0068, **Fig. 4e**). In contrast, there was no survival difference between male and female SG mice (*P* = 0.2, **Fig. 4f**). These results suggest that *Yap1* function is more critical for MB progression in males.

**Figure 4:**
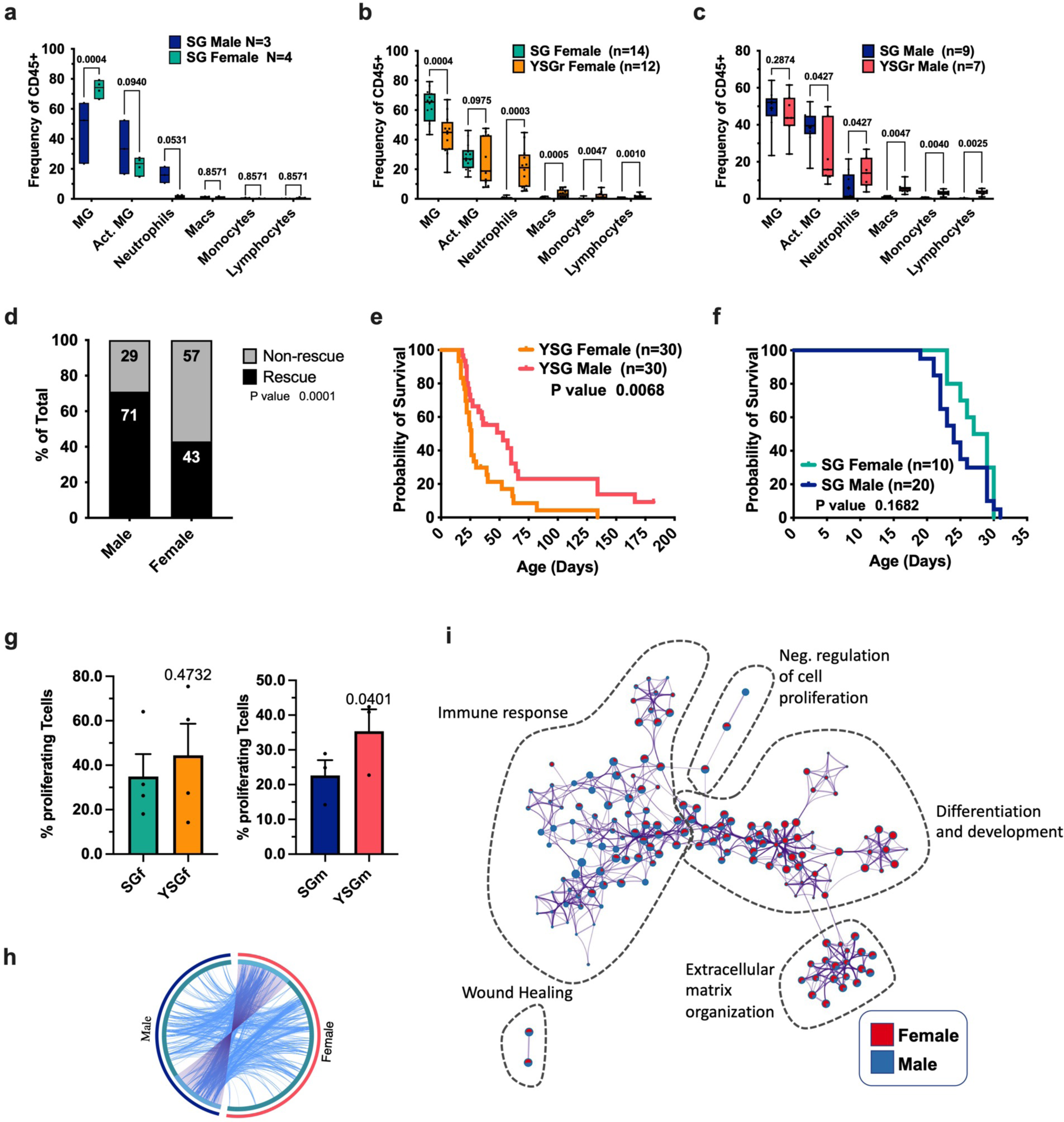
*Yap1* simultaneously promotes stemness and suppresses neural differentiation. **a-c.** Quantification of SG male vs female post-weaning age samples only (a), female SG vs YSG^r^ (b), or male SG vs YSG^r^ (c) by flow cytometry analysis. Number of mice is indicated in graph legend. Line inside box indicates median values. *P-*values were calculated using two-way ANOVA tests followed by the two-stage linear step-up procedure of Benjamini, Krieger, and Yekutieli. **d.** *Yap1* deletion rescues MB formation and extends survival in *Fsmo;GFAP-cre* mice in a sex-biased manner. Contingency analysis shows significant male bias in YSG^r^ tumors. *P*-value represents chi-squared test. **e.** Kaplan-Meier survival curves of male and female YSG mice. **f.** Kaplan-Meier survival curves of male and female SG mice. **g**. T cell proliferation assay using dye dilution. T cells were isolated from SG spleens, labeled, and cocultured sex-matched with SG or YSG cells for 4 days. *P*-value from paired Student’s *t-*tests. Whiskers represent minimum and maximum values, the line inside the box represents the mean, and the box extends from the 25th to 75th percentiles. n = 3 each (male) and n = 4 each (female) SG spleens. **h-i.** Metascape analysis of DE genes between YSG and SG male and female samples. Circos plot (h) showing overlapping DE genes between male YSG vs SG and female YSG vs SG samples. Purple lines represent shared genes, and blue lines represent genes sharing GO annotation terms (LFC >1, FDR <0.05). Gene Ontology (GO) network (i), each term is represented by a pie chart. Each pie sector is proportional to the number of hits originated from either male or female DE gene lists (please refer to Supplementary Fig. 2c for more insights).

To functionally compare *Yap1* expression in male and female MB cells, we isolated T cells from SG mouse spleens and cocultured them with sex-matched SG or YSG cells. Male YSG MB cell cocultures showed significantly higher T cell proliferation compared with male SG MB cells (**Fig. 4g**, *P* = 0.0401). In contrast, there was no significant difference between female YSG and SG MB cells in stimulating T cell proliferation (**Fig. 4g**, *P* = 0.4732). These data strongly support our survival analysis and indicate that *Yap1* function is sex-biased and more critical in male MBs.

To test this surprising finding in an unbiased manner, we performed Metascape multi-gene-list analysis^58^ of top genes differentially expressed (DE) between YSG and SG male (N=358) and female (N=307) samples (fold-change >2, FDR < 0.05). Of these, only 84 genes overlapped (**Fig. 4h**), indicating a sex-specific transcriptional program downstream of *Yap1*. Gene regulatory network analysis using GO annotation terms, clustered by similarity and color labeled by number of genes contributing to the enrichment from either male or female gene lists, revealed a greater contribution of female DE genes in development and differentiation-related clusters (**Fig. 4i, Supplementary Fig. 6b**). On the other hand, male DE genes contributed highly to the immune response-related clusters (**Fig. 4i, Supplementary Fig. 6b)**.

To test whether expression of *Yap1* is sex biased in MB, we compared *Yap1* expression levels between SG MBs from male and female mice. *Yap1* expression was not significantly different between male and female SG samples in our bulk RNA-seq datasets (**Supplementary Fig. 6c**). Furthermore, *Yap1* levels were also equivalently reduced in both male and female YSG MBs compared with SG MBs (**Supplementary Fig. 6c**). In human datasets, male SHH MB patients had higher levels of *YAP1* than females (**Supplementary Fig. 6d**), but *YAP1* did not stratify patient survival in any of the four subtypes (**Supplementary Fig 6e**). Together, these results suggest that the sex-biased function of *Yap1* cannot be explained by differences in *Yap1* RNA expression levels.

### Yap1 simultaneously promotes stemness, suppresses neural differentiation, and inhibits immune infiltration

To better understand the cell-autonomous effect of *Yap1* loss on MB tumor cells, we performed single-cell RNA-seq (scRNA-seq) analysis of SG and YSG male and female samples. We analyzed the transcriptional profiles of 31,150 single cells passing QC from SG male, SG female, YSG^r^ male, and YSG^r^ female samples (**Supplementary Fig. 7**). Unsupervised clustering of all aggregated cells revealed 19 clusters with distinct gene expression patterns **(Supplementary Fig. 7a-e and Supplementary Table 4**). Using marker genes for different lineages (**Supplementary Fig. 7b**), we assigned each cluster as tumor (clusters c1-12, c15-16; *Smo* and *Eyfp* (expressed by *SMO-M2*^+^ cells)), immune cells (c18; *Ptprc*/*Cd45*), endothelial cells (c19; *Col3a1*, *Pecam1/Cd31*), astrocytes (c14; *Aqp4, Gfap*), or oligodendrocytes (c17; *Sox10, Olig2*) (**Supplementary Fig. 7a-e**).

Most cells in our dataset were tumor cells (30,395 cells). Unsupervised *de novo* clustering of MB cells only (**Supplementary Fig. 7a,c**) identified 10 subclusters (MC01-10) with unique gene expression patterns (**Supplementary Fig. 7f-h and Supplementary Table 5**). Marker gene and GSEA analyses showed that clusters MC01, MC05, and MC10 resembled cycling cGNPs; MC02, MC03, MC07, and MC09 resembled G1 cGNPs; and MC04, MC06, and MC08 resembled mature cGNPs (**Fig. 5a, Supplementary Fig. 7i,j**). SG male and female cells dominated clusters MC01, MCO3, and MC06 while YSG male and female cells dominated MC02, MC04, and MC05 clusters. Clusters MC07 and MC08 were mixed (**Supplementary Fig. 7f, g, h**). An overall comparison of DE gene sets between samples showed reduced proliferation and stem cell pathway signatures and an enriched neuronal pathway signature in both male and female YSG cells (**Fig. 5b**). These results are consistent with the *in vitro* self-renewal and histological analyses shown in **Fig. 1b-e** and suggest a shift in developmental trajectory from stem-like proliferating cells to more mature postmitotic cells in YSG MBs.

**Figure 5:**
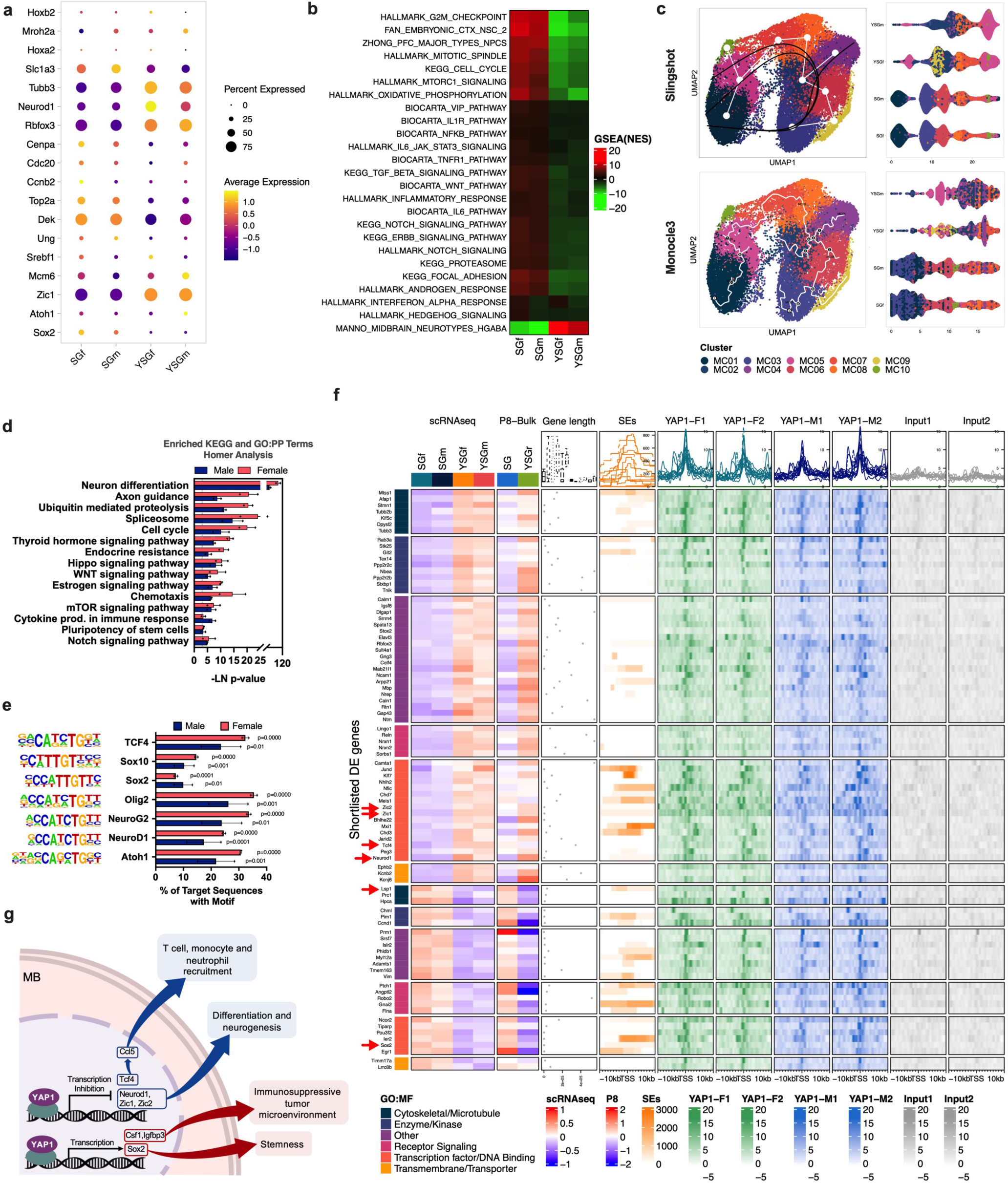
YAP1 simultaneously promotes stemness and suppresses neural differentiation. **a.** Dot plot showing the expression of neuronal markers in tumor clusters only. Dot sizes indicate the percentage of cells in each sample expressing the gene, and colors indicate average expression. **b.** GSEA analysis of differentially enriched Hallmark and C8 pathways showing enrichment of mature neuron signatures in YSG samples compared with SG. **c.** Pseudotime analysis using Slingshot or monocle3 confirming increased differentiation status in YSG samples compared with SG samples. **d-f**. Analysis of YAP1 ChIP-seq from 2 independent batches, each batch performed with 3 pooled male or female SG samples. **d.** KEGG and GO:PB pathway enrichment analysis of YAP1 binding regions using Homer. **e.** Homer motif enrichment analysis showing overrepresented TF binding sites within YAP1 peaks. Error bars represent SEM and dots represent data from each batch. **f.** EnrichedHeatmap showing common DE genes in scRNA-seq (row-centered LogCounts), Bulk RNA-seq(p8-row-centered Log2CPM) annotated by their most frequent GO:MF term and comparing the gene length with super enhancer position and tag distributions of YAP1 ChIP in female or male SG samples around transcription start sites (TSS; +10/-5 kb) presented both as average plots (average of values for all target regions in y-axis-top) and as heatmaps (values on z-axis as color, regions on y-axis - bottom ). The red arrows highlight genes of interest. **g.** Schematic representation of the proposed mechanism.

To test this hypothesis, we performed pseudotime trajectory analyses using Monocle3^59^ and Slingshot^60^ R packages (**Fig. 5c**). Both trajectory algorithms showed cascading developmental trajectories from proliferating stem/early NSC-like cells to late/mature cerebellar granule neuron-like cells, consistent with marker expression in these populations (*Sox2àAtoh1àNeuroD2/Cntn1*) (**Supplementary Fig. 8**). Notably, YSG cells were dramatically reduced in stem/progenitor points and increased in late/mature cGNP-like states. YSG MB cells showed loss of both cycling and G1 *Top2a/Cenpa/Sox2*^+^ stem-like/early cGNP-like cells and an increase in *Neurod1/Cntn1/Rbfox3/Tubb3*^+^ late cGNP/differentiating neurons (**Fig. 5a, Supplementary Fig. 7f, Supplementary Fig. 8a,b**). These results strongly indicate that *Yap1* promotes stemness in MB cells in a cell-autonomous manner and provides a molecular mechanism underlying the partially normalized cerebral development in rescued YSG tumors (**Fig. 1b**).

Single cell analysis of YSG MB cells also showed reduced enrichment for gene sets associated with inflammatory responses including IL6-JAK-STAT3, TNFR1, NFKB, IL1R, and TGFβ signaling (**Fig. 5b**), supporting a cell-autonomous regulation of immune response in MB cells by *Yap1 in vivo*.

To determine whether YAP1 regulates these pathways directly as a transcription factor, we performed YAP1 ChIP-seq to identify targets downstream of YAP1 in MBs *in vivo*. Chromatin preparations from male and female SG MB tissues were used to pull down YAP1-binding sites *in vivo*. 34.6% of female and 42.7% of male peaks were located at promoter regions, 28% of female and 22.4% of male peaks were located at intronic regions, and 25.2% of female and 18.7% of male peaks were located at intergenic regions (**Supplementary Fig. 9a, Supplementary Tables 6-9**). Differential peak calling analysis showed 14 significant differences in YAP1-binding sites in male vs. female SG samples (**Supplementary Fig. 9b,c; Supplementary Table 10**). The 14 differentially enriched YAP1 binding sites in the female vs. male samples (FDR <0.05, fold-change >2) were all located on the X chromosome and encoded 4 ncRNAs [*Xist* (X chromosome), *Firre* (X chromosome), *4933407K13Rik* (*Dxz4*) (X chromosome), and *Gm10494* (chromosome 17)] (**Supplementary Fig. 9b-d, Supplementary Table 10**). At the expression level, two of three female YSGr MBs showed lower RNA levels of *Xist, Firre*, and *4933407K13Rik* (*Dxz4*) in our bulk RNA-seq endpoint dataset compared with SG MBs (**Supplementary Fig. 9e**). However, the averaged expression of these genes was not significantly different in female YSG vs. SG MBs (**Supplementary Fig. 9e)**, suggesting Yap1 is not a major regulator of their expression.

KEGG and GO pathway enrichment analyses of YAP1-binding regions showed that YAP1 binding sites are highly enriched for genes associated with neuron differentiation, cell cycle, Hippo signaling, WNT signaling, chemotaxis, and cytokine production in MBs (**Fig. 5d**). Notably, Homer^61^ motif enrichment analysis revealed that YAP1-bound regions overlapped with SOX10, SOX2, NEUROD1, NEUROG2, ATOH1, and OLIG2 binding motifs (**Fig. 5e, Supplementary Table 11**). These are key transcriptional regulators of cerebellar neurogenesis, suggesting that YAP1 may cooperate or compete with these transcription factors to regulate neural stem cell and progenitor fates in the cerebellum. Unexpectedly, TEAD binding sites were not among the top 30 enriched motifs (**Supplementary Table 11),** suggesting that YAP1 binds to other partners in MB cells to promote stemness and inhibition differentiation.

To identify targets downstream of YAP1 in SHH MBs, we integrated ChIP-seq data with scRNA-seq (MB cells only) and bulk P8 RNA-seq (**Fig. 5f**) datasets to identify genes associated with YAP1 binding and whose expression levels are altered in YSG samples. The intersection of the differentially expressed gene lists between YSG and SG tumors and YAP1-binding sites represented the highest confidence YAP1 direct targets: 89 genes with YAP1 binding were down- or upregulated upon *Yap1* loss in YSG MB cells during the early stage of tumor initiation (p8 data) and in end-stage MBs (endpoint scRNA-seq) (**Fig. 5f**). 45 out of 89 genes had YAP1 binding in super-enhancer regions (**Fig. 5f**), consistent with a previous report of YAP1 binding in super-enhancers^32^ . Notably, 22 out of 89 genes encoded transcription factors/DNA binding proteins (*P* = 0.003), 39 genes were involved in nervous system development (*P* = <0.000001), and 29 were involved in neuronal differentiation (*P* = <0.000001).

YAP1 binds to the promoter regions of key cerebellar neural stem and progenitor regulators such as *Sox2*, *Neurod1*, *Zic1*, and *Zic2* (**Supplementary Fig. 9f)** and regulates their transcription. These transcription factors were differentially expressed in both our scRNA-seq and bulk RNA-seq datasets, indicating that *Yap1* promotes stemness through direct transcriptional regulation of these genes (**Supplementary Fig. 9g,h**). Further, gProfiler (g:GOSt)^62^ TRANSFAC database analysis of DE genes between YSGr and SG tumor cells in our scRNA-seq dataset (392 genes, FDR <0.05, logFC >0.25) showed that 25.5% of DE genes contained NEUROD transcription factor binding site (TFBS; Motif: NNSCWGCTGNSY, adj *P-*value = 1.47E-19), 24.7% contained SOX2 TFBS (Motif: CCWTTGTYATGCAAA, adj *P*-value = 6.10E-13), 54.1% contained ZIC1 TFBS (Motif: KGGGTGGTC, adj *P-*value = 2.30E-50), and 25.5% contained ZIC2 TFBS (Motif: NNCCCCCGGGGGGG, adj *P-*value = 1.87E-19). Taken together, our data strongly suggest that YAP1 simultaneously promotes stemness through the activation of *Sox2* transcription and inhibition of pro-differentiation factors *Neurod1* and *Zic1/2*, and we propose a transcriptional regulatory network outlined in **Fig. 5g**.

Next, we analyzed whether the cytokines/chemokines shown in **Fig. 3h** are direct YAP1 targets. YAP1 binds to the genomic regions of *Csf1*, *Igfbp3*, and *Il1* (**Supplementary Fig. 9i**). Further, *Csf1* and *Igfbp3* expression levels were reduced in both male and female YSG tumor cells compared with SG cells by scRNA-seq analysis (**Supplementary Fig. 9j**). Rantes/Ccl5 expression was highly elevated in YSG MB cells compared with SG cells (**Supplementary Fig. 9j**), potentially explaining increased TIL infiltration in YSG MBs. However, YAP1 does not bind to the *Ccl5* genomic region (**Supplementary Fig. 9i**), suggesting that Yap1 modulation of *Ccl5* expression is indirect. Accordingly, we searched the CHEA Transcription Factor Binding Site datasets^63^ to identify known regulators of *Ccl5* transcription. Expression analysis of ten potential transcription regulators of Ccl5 expression in our scRNA-seq dataset (**Supplementary Fig. 9k**) showed that *Tcf4 (*CHEA::*TCF4-23295773-U87-HUMAN)* expression was significantly upregulated in YSG vs. SG MB cells (**Supplementary Fig. 9k**). *Tcf4* has multiple YAP1 peaks in super-enhancers near the *Tcf4* locus (**Supplementary Fig. 9l)**, suggesting that YAP1 regulates *Ccl5* expression indirectly by regulating *Tcf4* expression.

### Yap1 targets stratify male MB patient survival

To assess the relevance of direct YAP1 targets in human MB, we analyzed associations between YAP1 target gene signatures and survival in a human MB dataset (GSE85217)^26^. Univariate survival analysis of high-confidence YAP1 target genes showed that 31 predicted survival in human MB patients (**Supplementary Table 12**; only 80 of 89 target gene homologs were present in the GSE85217/Cavalli *et al.* dataset). Strikingly, when we grouped the genes by *Yap1* regulation (i.e., repressed vs. activated), we discovered that 26% of YAP1-repressed genes predicted a favorable outcome and 7% of YAP1-repressed genes predicted a poor outcome in male patients. In contrast, only 2% of YAP1-repressed genes predicted a good outcome and 4% YAP1-repressed genes predicted a poor outcome in female patients (**Fig. 6a**). Notably, a gene signature composed of the top 14 YAP-repressed genes predicted better survival in male patients but not female patients (**Fig. 6b,c**). In addition, 28% of YAP1-activated genes predicted a poor outcome and 8% of YAP1-activated genes predicted a good outcome in male patients. In contrast, only 4% of YAP1-activated genes predicted a poor outcome and 8% of YAP1-activated genes predicted a good outcome in female patients (**Fig. 6d**). A gene signature composed of the top 7 YAP-activated genes predicted poor survival only in male patients (**Fig. 6e-f**). These results highlight an evolutionarily conserved sex-biased function of YAP1 in MB and provide important new insights into sexual dimorphism in MB.

**Figure 6:**
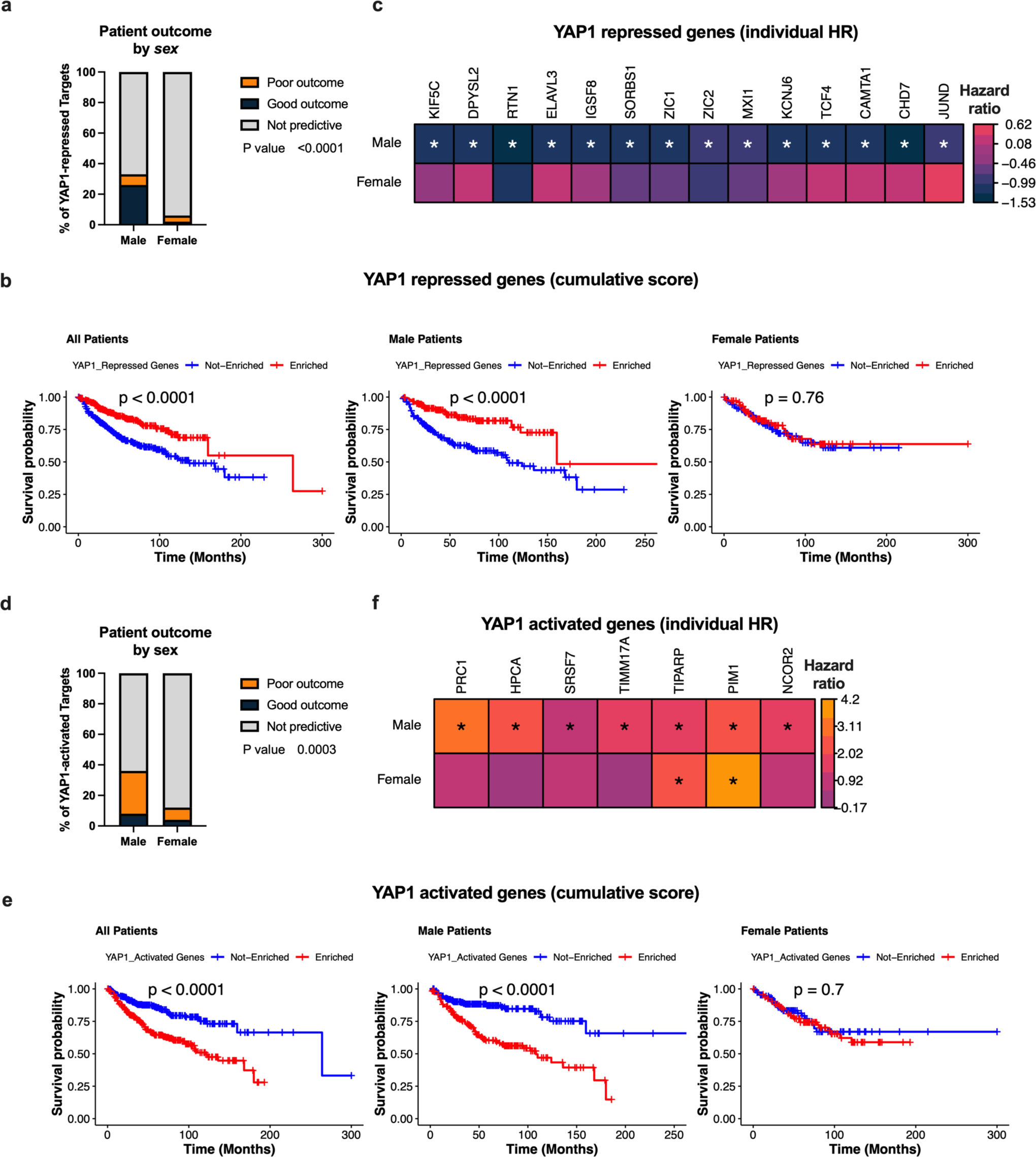
YAP1 targets stratify male MB patient survival. **a-c** Survival analyses of YAP1-repressed genes in the Cavalli *et al.* dataset. (a) Contingency analysis showing the frequency of YAP1-repressed targets predictive of patient survival stratified by sex. *P*-value represents chi-squared test. (b) A 14-gene signature composed of YAP1-repressed genes as a Kaplan-Meier plot or (c) individual YAP1-repressed genes as a heatmap. Heatmap shows hazard ratio of high expression. Color indicates scaled and centered hazard ratios as calculated by survival::coxph() (exp(coef)). Asterisks indicate Cox’s regression model *P*-values <0.05. **d-f.** Survival analyses of YAP1-activated in the Cavalli *et al.* dataset. (d) Contingency analysis showing the frequency of YAP1-activated targets predictive of patient survival stratified by sex. *P*-value represents chi-squared test. (e) A 10-gene signature composed of YAP1-activated genes as a Kaplan-Meier plot or (f) individual YAP1-activated genes as a heatmap. Heatmap shows hazard ratio of high expression. Color indicates scaled and centered hazard ratios as calculated by survival::coxph() (exp(coef)). Asterisks indicate Cox’s regression model *P*-values < 0.05.

## DISCUSSION

Elucidating the mechanisms underlying sexual dimorphism in cancer is an emerging area of intensive research^5, 64^. However, our understanding of the interplay between oncogenic events and sex as a variable in tumorigenesis, particularly in brain tumors, is still primitive. We discovered a sex-biased requirement for *Yap1* in SHH MB progression. *Yap1* deletion in male SG mice extended their survival more often and for a longer than its deletion in female SG mice. This is particularly interesting since, in humans, MB is more common in males at all ages^12, 13^. Additionally, non-infant males have a considerably worse outcome/survival than females^17, 18^.

Mechanistically, *Yap1* directly promoted stemness and blocked the differentiation of transformed cGNPs in SHH MBs by at least two distinct but complementary mechanisms. First, YAP1 directly activated the transcription of a key MB stem cell regulator, SOX2. Concurrently, YAP1 blocked maturation of MB stem cells by inhibiting the transcription of cerebellar neural stem and progenitor maturation regulators, *NeuroD1* and *ZIC1*/2. In this way, YAP1 expression generates a reinforced transcriptional regulatory network that promotes stem cell maintenance and suppresses differentiation. Second, YAP1 amplifies its effect by modulating the function of these key lineage determining transcription factors at the protein level. Unexpectedly, we discovered that the YAP1 binding sites in MB cells *in vivo* did not overlap with canonical TEAD binding sites; rather they were enriched for consensus binding sites for SOX2, ATOH1, NEUROD1, and ZIC1/2, the same transcriptional regulators of cerebellar NSC and cGNP fates controlled by YAP1 at the RNA level. In other words, YAP1 protein binds to the same target genomic regions bound by SOX2, ATOH1, NEUROD1, and ZIC1/2. We hypothesize that YAP1 cooperates with SOX2 to regulate its downstream genes that promote stemness and competes with NEUROD and ZIC1/2 for binding to the regulatory regions of their downstream genes that promote differentiation. In support of this hypothesis, ∼75% of all differentially expressed genes in SG vs. YSG MBs downstream of YAP1 had binding sites for SOX2, ATOH1, NEUROD1, and ZIC1/2 proteins that overlapped the YAP1 binding sites in their regulatory regions. Based on these results, we propose a novel molecular network in which a multi-level YAP1 function reinforces the maintenance of stemness and inhibition of differentiation in SHH MB.

The functional importance of SOX2 and YAP1 as MB stem cell regulators has been independently demonstrated in previous studies. Vanner *et al.* showed that SOX2^+^ cells in SHH MB are quiescent cancer stem cells^42^, and Fernandez *et al.* reported that YAP1^+^ perivascular MB cells survive radiation treatment and re-seed MB recurrence^27^. Our combined ChIP-seq and scRNA-seq analyses mechanistically link YAP1 and SOX2 and show that YAP1 is an upstream regulator of SOX2 in SHH MBs *in vivo*. Furthermore, we discovered that the major binding partners for YAP1 in SHH MBs *in vivo* are not TEAD proteins. There was no enrichment of TEAD binding sites in our dataset, even though we used a previously validated YAP1 antibody for ChIP-seq. Indeed, ChIP-seq using the same YAP1 antibody in cardiomyocytes pulled down TEAD binding sites^65^ (**Supplementary Table 13**), strongly suggesting that YAP1 binding partners vary significantly in different tissues and that YAP1 function is mediated through its binding partners in a context-dependent manner.

In terms of a sex-biased requirement for YAP1 in SHH MB, our data point to *Yap1* expression in MB cells having a sexually dimorphic effect on inflammatory response and STAT3 activation. Functionally, we observed that Yap1-deleted male MBs stimulate T cell proliferation which was not the case in female MB cells (**Figure 4g**). Consistently, our Metascape multi-gene-list analysis with differentially expressed genes between SG and YSG tumors show enrichment of immune response/inflammatory network in males more so than in females (**Figure 4i**). At the gene level, the only genes differentially regulated by YAP1 in male vs. female MBs were X-chromosome-associated non-coding RNAs *Xist*, *Firre*, and *Dxz4* (**Supplementary Fig. 9d,e**). *Xist* has previously been suggested to play a role in sex-biased cancer risk^66^. *Xist* normally silences one of the two X chromosomes in female cells, but recent studies have shown that X inactivation is imperfect and that there is leaky expression from the silenced X chromosome, elevating the expression of ∼10-20% of X chromosome genes^66^. This is notable, since the X chromosome is enriched for genes encoding cytokines/chemokines and other important immune modulatory genes such as *FoxP3* critical for Treg function and reported to be regulated YAP1. In addition, while it was originally believed that *Firre* and *Dxz4* are regulators of X chromosome inactivation, a recent publication suggests their involvement in autosomal gene regulation *in vivo*^67^. Together, these results suggest that *Yap1* loss in females, more so than in males, might be accompanied by further changes in gene expression downstream of *Firre* and *Dxz4* that may contribute to sex-biased rescue of MB formation in males.

Previous studies have reported both cell-autonomous and non-autonomous functions for *Yap1* in immune suppression. *YAP1* is essential for T-reg differentiation and function in a cell-autonomous manner^35^. Additionally, YAP1 regulates cytokine secretion by multiple cancer cell types including colorectal cancer^36^, prostate adenocarcinoma^37^, and pancreatic ductal carcinoma^41^ to recruit immune suppressive myeloid derived suppressor cells (MDSCs) and M2 macrophages^36, 37, 41^. We showed that *Yap1* deletion in MB cells significantly increased T cell and BMDM infiltration (including neutrophils, monocytes, and macrophages). This contrasts with YAP1 function reported in pancreatic and prostate cancer cells, where *Yap1* expression promote MDSC infiltration^37, 41^. Mechanistically, YAP1 directly activates transcription of *Csf1* and *Igfbp3,* which are known to promote immune suppression^68, 69^. In addition, *Yap1* expression in MB cells enhances IL6-STAT3 and TGFβ signaling (**Fig. 5c**), which also promotes an immune suppressive microenvironment. YAP1 also indirectly blocks the expression of *Ccl5* by regulating the expression of *Tcf4*: YAP1 does not bind to *Ccl5* regulatory regions but does bind to and regulate *Tcf4* expression, which in turn can regulate *Ccl5* expression. Since Ccl5 is known to recruit T cells^54^, monocytes^54^, and neutrophils^55^, we propose that YAP1 induces an immune suppressive tumor microenvironment in MB by directly activating transcription of *Csf1* and *Igfbp3* and indirectly suppressing *Ccl5* transcription via *Tcf4* (**Fig. 5g**). We also discovered that *Lsp1*, known to inhibit neutrophil and lymphocyte recruitment^70, 71^, is also a direct YAP1 target (**Fig. 5f**). Finally, our analysis shows that *Tcf4* and *Igfbp3* levels are altered in YSG SHH MB as early as postnatal day 8, suggesting that YAP1 expression in early-stage cancers may promote immune evasion.

A recent study highlighted a male-biased role for STAT3 in an SHH mouse model^64^. White *et al.* showed that conditional *Stat3* deletion in *Math1*-positive cells protected male *Ptch1*^LacZ/+^ mice from developing MBs. However, they reported that *Stat3* deletion has no effect on the proliferation, apoptosis, and differentiation of *Ptch1*^LacZ/+^ tumors^64^. In our model, single-cell RNA-seq and western blot analyses showed that both male and female YSG MBs have reduced *Stat3* RNA and protein levels in both male and female YSG MBs compared to SG MBs (**Supplementary Fig. 10a-c**). This suggests that YAP1 is upstream of STAT3 in SHH MB and that reduced STAT3 levels may also contribute to male bias and stem cell regulation.

Interestingly, we discovered that STAT3 phosphorylation is sexually dimorphic (**Supplementary Fig. 10c**). While phosphorylation of STAT3 at Tyr705 was not significantly different between SG and YSG MBs in both males and females, pSTAT3 at Ser727 was significantly higher in male YSGs but not female YSGs. A previous study showed that pSTAT3 Y705 is required for STAT3-mediated self-renewal of mouse embryonic stem cells, while pSTAT3 Ser727 phosphorylation is crucial for transitions from pluripotent stem cells to neuronal commitment^72^. Together, these results suggest that in male SHH MB, phosphorylation of STAT3 at S727 is downstream of YAP1 and may contribute to suppressed neuronal differentiation. A recent study in gliomas showed a significant correlation between YAP1 and pSTAT3-S727^73^, supporting our model that partially explains the underlying mechanism of male-biased YAP1 function.

Emerging studies suggest that immune checkpoint inhibitors, specifically anti-CTLA4, significantly improve overall survival and progression-free survival in male cancer patients more so than in female patients^74^. This supports sex-dependent dimorphism in the tumor microenvironment and immune suppression^75^. MBs are considered immune “desert” tumors, since they have a highly immune suppressive microenvironment with few infiltrating lymphocytes^24, 76, 77^. Immune checkpoint inhibitors, such as anti-PD-1 or anti-PD-L1 blockade, are of only limited benefit since there are very few infiltrating T cells and PD-L1 expression is low in MBs^24, 76, 77^. In addition, SHH MBs have greater numbers of infiltrating MDSCs and fewer PD1^+^ T cells than group 3 MBs ^78–80^. This study strongly suggests that YAP1 is a major culprit responsible for immune suppressive TME and male biased tumorigenesis in MB.

In conclusion, our *in vivo* and *in vitro* studies strongly suggest that *Yap1* is a significant cooperating oncogene in SHH-induced MB progression, and it functions as a critical regulator of SHH MB CSCs and immune evasion. Notably, YAP1 function is more critical in males than females, and this is evolutionarily conserved: YAP1 downstream target genes we identified are significant stratifiers for survival in male but not female MB patients. These results nominate a set of novel genes for future investigation to elucidate additional underlying causes of sexually dimorphic incidence and prognosis of MB. In addition, our results strongly suggest that male MB patients will likely benefit from YAP1 inhibitors more than female MB patients. This study provides compelling evidence for incorporating sex as a major variable in mechanistic and therapy response studies involving YAP1 as an oncogene in all cancers in which the Hippo/YAP1 pathway is mutated or activated.

## MATERIALS AND METHODS

### Mouse models of SHH MB

The following mouse models were generated or used for use in this study:*Ptch1^+/-^* mice (Stock: Ptch1^tm1Mps^/J, RRID:IMSR_JAX:003081), *Trp53^+/+^* mice (Stock: B6.129S2-Trp^53tm1Tyj^/J, RRID:IMSR_JAX:002101), *FSMO-M2* mice: (Stock: Gt(ROSA)26Sort^m1(Smo/EYFP)Amc^/J, RRID:IMSR_JAX:005130), h*GFAP*^cre^ mice: (Stock: FVB-Tg(GFAP-cre)25Mes/J, RRID:IMSR_JAX:004600), *Yap1*^fl/fl^ mice were from Dr. Fernando Camargo^81^, *Olig2*^Cre^ mice (stock: B6.129-Olig2^tm1.1(cre)Wdr^/J, RRID:IMSR_JAX:025567), *NeuroD2-SmoA* (stock: C57BL/6-Tg(Neurod2-Smo*A1)199Jols/J, RRID:IMSR_JAX:008831) Both male and female mice were utilized in this study. Mice were randomly assigned to each group and were age and sex matched at the time all experiments. Mice were housed and handled in accordance with the protocols and procedures approved by the HMRI and The Jackson Laboratory Institutional Animal Care and Usage Committees.

### Primary cell lines and cell culture

Primary tumorsphere cells were established from *Ptch;p53* or *FsmoM2;GFAP-Cre* tumors by cutting micro-dissected tissues into small chunks and then treating them with Accutase™ (Corning Inc., Corning, NY; #25058CI) to generate single cell suspensions. These were cultured in standard Neural Stem Cells (NSC) medium (DMEM/F12 (HyClone, Logan, UT; #SH30261.01), B-27™ supplement (1:1000, Thermo Fisher Scientific, Waltham, MA; #17504044), penicillin-streptomycin (GenDEPOT, Katy, TX; #CA005-010), 10 ng/ml bFGF (Peprotech, London, UK; #450-33), and 20 ng/ml EGF (Peprotech; #315-09)). For secondary sphere formation assays, 3000 viable, single cells were plated into six-well plates with 3 mls of medium and counted 6-7 days later. For *Yap1* siRNA treatment; *Ptch;p53* primary cell-lines (#9410) were transfected with 40 nM ON-TARGETplus mouse *Yap1* siRNA (GE Healthcare Dharmacon, Inc., Lafayette, CO; # L-046247-01-0005) using Lipofectamine™ RNAiMAX (Invitrogen™, Waltham, MA; #13778075). To develop CRISPR mediated KO of *Yap1*, *Ptch;p53* primary cell-line(#4954) was transduced using viral particles containing the lentiCas9-Blast vector (Addgene, Watertown, MA; RRID:Addgene_52962). Cells stably expressing Cas9 were selected using 10ug/ml Blasticidin S HCl (Gibco™-#R21001). 4954-Cas9 cells were then transfected with 30µM control sygRNA (Millipore Sigma, Burlington, MA; NegativeControl1, CGCGAUAGCGCGAAUAUAUU), *Yap1* sygRNA#1 (Millipore; MMPD0000043027, GCCGUCAUGAACCCCAAGA), *Yap1* sygRNA#2 (Millipore; MMPD0000043029, CGAAAGCAUGGCCUUCCGG) using Lipofectamine™ RNAiMAX (Invitrogen™; #13778075). We clonally selected 3 lines from each guide RNA for further study.

### Intracranial injections

Freshly dissociated PDX tumor cells were injected into the cerebella of male and female 6-8 weeks old NSG mice (RRID:IMSR_JAX:005557) using a stereotaxic device (Bregma: 1/-6.5/-2.8). To eliminate potential spatial heterogeneity within the parental tumor, each tumor was first minced into slurry and equivalent pools of tumor cells were injected. The following PDXs were used in this study: Med1712 (SHH subtype, Source: James Olson-Brain Tumor Resource Lab) and Med211 (Group3 subtype, source: James Olson, Brain Tumor Resource Lab)*. Ptch;p53* primary cell-line(#4954) Cas9– control, sygRNA 1.2 or sygRNA 2.2 were injected into the cerebella of male and female 6-8 weeks old B6 mice (RRID:IMSR_JAX:000664) using a stereotaxic device (bregma: 1/-6.5/-2.8). All procedures were approved by the HMRI and The Jackson Laboratory Institutional Animal Care and Usage Committees.

### Immune phenotyping by flow cytometry

Freshly dissected tissue was micro-dissected into small chunks and then treated with Accutase™ for 10-15 min at 37°C. Accutase™ was removed and tissues were resuspended in complete medium to triturate and generate single-cell suspensions. Cells were resuspended in RBC lysis buffer to remove red blood cells. Following RBC lysis, cells were strained through 40µm Flowmi cell strainers (Bel-Art, Wayne, NJ; #H136800040). Cells were then stained with multiple flow cytometry validated antibody cocktails (see below) and analyzed using either BD Fortessa or LSRII cytometers (BD Horizon, BD, Franklin Lakes, NJ). Data were analyzed and quantified using *FlowJo* v10.6 software (RRID:SCR_008520). All antibody dilutions and staining were performed in Brilliant Stain Buffer (BD; #563794), and cells were incubated with a blocking solution containing mouse TruStain FcX (Biolegend, San Diego, CA; #101319) before antibody staining. Antibodies used : CD45-PE-cy7 (Biolegend; # 103114, RRID:AB_312979), CD45-APC-cy7 (Biolegend, San Diego, CA; # 103116, RRID:AB_312981), CD11b/Mac1-BV650 (Biolegend; # 101259, RRID:AB_2566568), CD3e-PE (Biolegend; # 100308, RRID:AB_312673), CD4-APC (Biolegend; # 100411, RRID:AB_312696), CD8-FITC (Biolegend; # 100706, RRID:AB_312745), Ly6c-BV711 (Biolegend; # 128037, RRID:AB_2562630), Ly6g-APC-cy7 (Biolegend; # 127624, RRID:AB_10640819)

### Immunofluorescence analysis

Tissues were fixed in 4% Paraformaldehyde (PFA) overnight, equilibrated through 10%, 20%, and 30% sucrose gradients, and then embedded in O.C.T compound (Fisher Healthcare, Thermo Fisher Scientific; # 23-730-571). Frozen samples were sectioned into 10μm thickness and slides were blocked in 5% normal goat serum/0.2%Triton PBS for 30min and incubated with primary antibodies (see below) overnight at 4 °C. Then, slides were incubated with appropriate Alexa Fluor secondary antibodies (Invitrogen™) for 30-45 min. Nuclei were stained with DAPI (1:2,000, Invitrogen™). The TrueVIEW autofluorescence quenching kit was applied (Vector Laboratories, Newark, CA; #SP-8500-15) to remove background fluorescence. Images were obtained using a Zeiss Axiovert 200M fluorescence microscope and the FV3000 confocal microscope (Olympus, Tokyo, Japan). Primary antibodies used : CD3e (Thermo Fisher Scientific Cat# 14-0032-85, RRID:AB_467054), CD11b (BD Biosciences Cat# 553308, RRID:AB_394772) ), IBA1 (Thermo Fisher Scientific Cat# PA5-27436, RRID:AB_2544912), CD45 (Millipore Sigma, Burlington, MA; #CBL1326, RRID:AB_2174425), SOX2 (R&D Systems, Minneapolis, MN; Cat# MAB2018, RRID:AB_358009) and Ki67 (Abcam, Cambridge, UK; Cat# ab15580, RRID:AB_443209).

### Immunohistochemical analysis

Tumor tissues were fixed in 10% formalin and embedded in paraffin. Paraffin blocks were sectioned to 5 μm thickness, deparaffinized and boiled in 10mM sodium citrate (pH 6.0) buffer to retrieve antigens. Slides were blocked in 5% goat serum (Sigma Aldrich, St Louis, MO; #G9023) for 30 min and incubated with YAP1 (Proteintech Cat# 13584-1-AP, RRID:AB_2218915) or KI67 (Abcam Cat# ab15580, RRID:AB_443209) primary antibodies overnight at 4°C. Then, slides were washed and incubated with anti-mouse/rabbit/goat IgG-biotinylated secondary antibody (Vector Laboratories; # BA-1300, RRID:AB_2336188) for 30-60 min, followed by ABC (Vector Laboratories; #PK-6100) for 1 hr and DAB (Vector Laboratories; # SK-4105) chemogenic reaction. Nuclei were counter stained with Hematoxylin ( VMR, US; #95057-844). Images were captured using an Olympus BX 41 microscope (Olympus).

### Western blotting

Tissues were lysed in NP-40 buffer (GenDEPOT) supplemented with protease and phosphatase inhibitor cocktail (GenDEPOT). Lysates were quantified using the DC Protein Assay (BioRad, Hercules, CA). 40 µg of lysate was run on lab-made 10% SDS-PAGE gels. Gels were transferred onto PVDF membranes at 100 V, 4°C, for 40 minutes. Blots were washed in 1X TBS with 1% Tween-20 (TBST) and blocked with 1% BSA/TBST for one hour at room temperature. The blots were incubated with primary antibodies (1:1,000) in 1%BSA in TBST overnight at 40°C. The blots were washed in 1X TBST, incubated with secondary antibody (1:5,000) in 1%BSA/TBST for two hours at room temperature, and developed with ECL Immobilon Forte (Millipore Sigma; WBLUF0500). Primary antibodies were: phospho-STAT3 (Tyr705) (Cell Signaling Technology; # 9145, RRID:AB_2491009 ), phospho-STAT3 (Ser727) (Invitrogen; # 44-384G, RRID:AB_2533645), STAT3 ((Cell Signaling Technology; # 9139, RRID:AB_331757) ), ß-actin HRP Conjugated (GenScript; # A00730, RRID:AB_914100). Secondary Antibodies were: anti-mouse IgG HRP-(Cell Signaling Technology; # 7076, RRID:AB_330924) and anti-rabbit IgG HRP-linked (Cell Signaling Technology; # 7074, RRID:AB_2099233).

### Cytokine arrays and CCL5 ELISA

Secreted cytokine and chemokine profiles were determined in the cell culture supernatants of *Ptch;p53* cell-lines using commercially available protein arrays (RayBiotech Peachtree Corners, GA;mouse cytokine antibody array 3 and mouse cytokine antibody array 4) according to the manufactureŕs instructions. The absolute concentration. Of CCL5 was quantified using a commercially available ELISA kit (R& D R and D Systems;#DY478-05) in SG or YSG tumor lysates according to the manufactureŕs instructions.

### RNA sequencing analysis

Bulk RNA-seq analysis was carried out using the in-house pipeline at The Jackson Laboratory. *Trimmomatic* (version 0.33) was used to remove adapters and also leading and trailing low quality bases. Reads fewer than 36 bases long were discarded. Reads with more than 50% low-quality bases overall were filtered out, and the remaining high-quality reads were then used for expression estimation. Alignment estimation of gene expression levels using the EM algorithm for paired-end read data was performed using *RSEM* (v1.2.12 RRID:SCR_013027). RSEM uses *bowtie2* (RRID:SCR_016368) as an aligner to align the mapped reads against the mm10 reference genome. Data quality control was performed using *Picard* (v1.95-RRID:SCR_006525) and *bamtools* (RRID:SCR_015987) to obtain general alignment statistics from the BAM file. The read counts estimated for each gene by RSEM were given as input to the R package *edgeR* (v3.30.3-RRID:SCR_012802) for differential expression analysis.

### Pathway analysis

Gene Set Enrichment Analysis (GSEA). We used *fGSEA* (v1.14.0 - RRID:SCR_020938) R package to test for enrichment of the Hallmark genesets downloaded from MsigDB (RRID:SCR_016863, msigdbr R package v7.2.1). For input, we used either z-score statistics from *EdgeR* (bulk RNAseq) DE analysis or pre-ranked gene lists generated using a fast Wilcoxon rank-sum test (scRNAseq) (*presto* R package v1.0.0 “github.com/immunogenomics/presto”). Gene Ontology Enrichment Analysis (GO). To identify enriched molecular pathways based on differentially expressed genes (DE genes), over-representation analysis (ORA) was performed on DE genes from each cluster using *g:Profiler* v0.2.0 (RRID:SCR_006809). We also used Ingenuity Pathway Analysis (IPA-Qiagen) and Metascape^58^ to provide more mechanistic insights.

### Quantitative Real-time PCR

RNA was extracted from tissues and cells with Trizol and treated with DNase (Ambion Inc., Austin, TX). cDNA was synthesized using iScript kit (BioRad) and real-time RT-PCR analyses were performed in triplicates using the iQ5 Optical System from Bio-Rad with SYBR Green mix (Life Technologies, Carlsbad, CA; # 4472953).18S was used as an internal control and relative fold change was calculated compared to normal cerebellar tissue or wildtype cerebellar NSCs. Primers used are listed below.

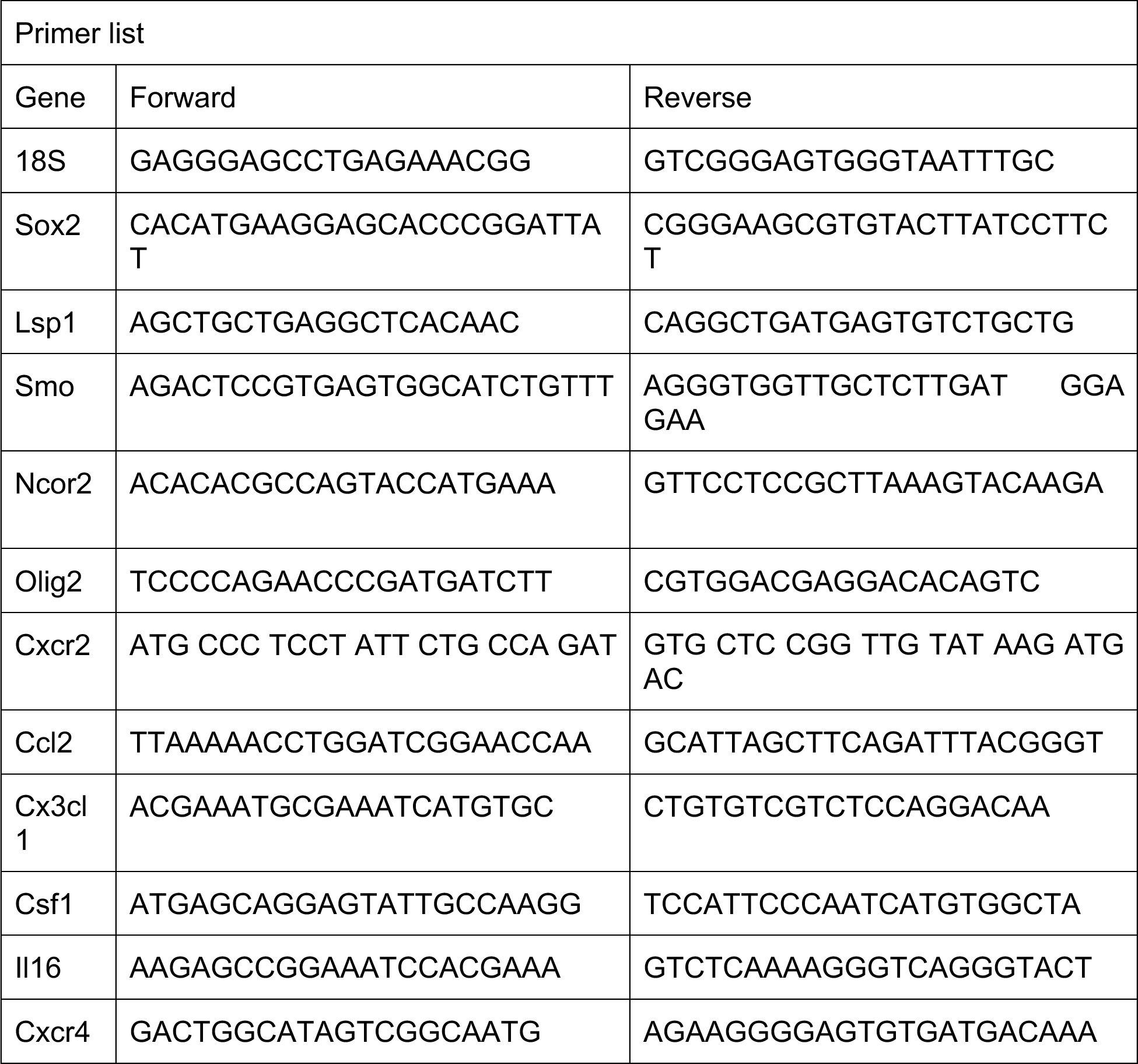

### Single cell RNAseq data collection

Freshly dissected tissue was micro-dissected into small chunks and then treating with Accutase™ for 10-15 min at 37°C. Accutase™ was removed, and tissues were resuspended in DME/F12+B27+pen/strep medium to triturate and generate single-cell suspensions. Cells were resuspended in RBC lysis buffer to remove red blood cells. Single cells were filtered using a 40µM nylon mesh (Falcon-#352340) to remove residual clumps. Dead cells were removed using dead cell removal kit (Miltenyi Biotec, Bergisch Gladbach, Germany; # 130-090-101). Cells were then counted and loaded onto the 10X Chromium controller, targeting 6000 cells for capture per well. Single-cell RNA sequencing libraries were generated using the Chromium Single Cell 3’ Library & Gel Bead Kit V1 or V3 and Chromium Single Cell 3’ Chips according to the manufacturer’s instructions (10X Genomics, Pleasanton, CA). In brief, All the single-cell samples and required reagents were loaded on a 10X Chromium controller for droplet generation, followed by reverse transcription in the droplets, cDNA amplification, fragmentation, adapter, and index addition following the manufacturer’s instructions. Barcoded single-cell transcriptome libraries were sequenced with 100bp paired-end reads on BGI’s DNBseq platforms.

### Single-cell RNA sequencing analysis

Raw sequencing reads from Illumina were aligned to mm10 genome (mouse) using *Cell Ranger* (v5; RRID:SCR_017344) software from 10X Genomics with default parameters. Subsequently, genes were quantified as UMI counts using *Cell Ranger* (v5;RRID:SCR_017344). Downstream analysis was performed on filtered feature-counts, generated by *Cell Ranger* and low-quality single cells containing < 200 expressed genes or > 20% mitochondrial transcripts were removed. Following the removal of low-quality cells, single cells were normalized and clustered using *Seurat*^82^ (v4.0.0; RRID:SCR_016341). Single-cell gene expression counts were normalized to the library size and log2-transformed. *Seurat*^82^ package was used to perform PCA reduction using the top 2000 most variable genes in the dataset. *Seurat*^82^ package was also used to identify cluster-specific marker genes and visualize dotplots and feature plots. The genes specifically expressed in each cluster were examined to identify the cell types. We used *ggplot2* (v3.3.3; RRID:SCR_014601) and *ComplexHeatmap* (v2.7.8.100; RRID:SCR_017270) packages in R to produce boxplots and heatmaps.

### Meta-analysis of an external human medulloblastoma scRNAseq dataset

We obtained scRNA-seq data from the Hovestadt *et al*.^50^ 10X single-cell RNA-seq dataset (#GSE119926, GEO - https://www.ncbi.nlm.nih.gov/geo/query/acc.cgi?acc= GSE119926). Count data were downloaded from GEO, and *Seurat*^82^ (v4.0.0; RRID:SCR_016341)was used to analyze the dataset as described above.

### YAP1 ChIP-seq analysis

Flash frozen fSMO;GFAP-cre MB tissues from males and females were sent to Active Motif (Carlsbad, CA) for ChIP-Seq analysis, using a previously ChIP validated anti-YAP1 antibody (US Biological, Salem, MA; # Y1200-01D-RRID:AB_2927438). Active Motif prepared chromatin, performed ChIP reactions, generated and sequenced the libraries, and performed basic data analysis. In brief, brain tissue samples were submersed in PBS + 1% formaldehyde, cut into small pieces, and incubated at room temperature for 15 minutes. Fixation was stopped by the addition of 0.125 M glycine (final). The tissue pieces were then treated with a TissueTearer and finally spun down and washed 2x in PBS. Chromatin was isolated by adding lysis buffer, followed by disruption with a Dounce homogenizer. Lysates were sonicated and the DNA sheared to an average length of 300-500 bp with Active Motif’s EpiShear probe sonicator (#53051). Genomic DNA (Input) was prepared by treating aliquots of chromatin with RNase, proteinase K and heat for de-crosslinking, followed by SPRI beads clean up (Beckman Coulter, Brea, CA) and quantitation by Clariostar (BMG Labtech, Aylesbury, UK). An aliquot of chromatin (200 µg) was precleared with protein A agarose beads (Invitrogen™). Genomic DNA regions of interest were isolated using 20 µg of antibody against YAP1 (US Biologicals-#Y1200-01D-RRID:AB_2927438). Complexes were washed, eluted from the beads with SDS buffer, and subjected to RNase and proteinase K treatment. Crosslinks were reversed by incubation overnight at 65 ℃, and ChIP DNA was purified by phenol-chloroform extraction and ethanol precipitation. Illumina sequencing libraries were prepared from the ChIP and Input DNAs by the standard consecutive enzymatic steps of end-polishing, dA-addition, and adaptor ligation. Steps were performed on an automated system (Apollo 342, Wafergen Biosystems/Takara). After a final PCR amplification step, the resulting DNA libraries were quantified and sequenced on Illumina’s NextSeq 500 instrument. The 75-nt single-end (SE75) sequence reads were mapped to the mouse mm9 genome using the *BWA* algorithm (RRID:SCR_010910) with default settings. Duplicate reads were removed, and only uniquely mapped reads (mapping quality ≥25) were used for further analysis. Known motifs were identified with the findMotifsGenome program in *HOMER* (v4.11; RRID:SCR_010881) using default parameters. Peak annotation and GO enrichment were performed using the annotatePeaks program in *HOMER*. BigWig files generated from BAM files using the bamCoverage function of *deepTools* (v3.5.0; RRID:SCR_01636) were used as input for *HOMER* analysis and visualization. Tracks shown in supplementary figure 9 were generated using the *trackViewer* R package (v1.24.2) and the heatmap shown in Figure 5f was generated using the *EnrichedHeatmap* R package (v1.18.2). *DiffBind* (v2.16.2; RRID:SCR_012918) R package was used for differential peak calling.

### Survival analysis and survival correlation

To assess the correlation between YAP target genes and survival of human medulloblastoma patients we used the publicly available human medulloblastoma dataset (Cavalli *et al*. ^26^; GEO Accession GSE85217). The combined signature scores of 10 up-regulated (YAP-repressed) or 7 down-regulated (YAP-activated) genes were generated using the score() function of the *JLaffy/scrabble* R package^83^ and survival analysis was performed using the *survival* (v3.2-7; RRID:SCR_021137) and *survminer* (v0.4.9; RRID:SCR_021094) R packages. Signature scores were centered, and all patient, male patients or female patients’ cohorts were stratified into two groups based on the sign of the signature score (above zero= “enriched”, below zero= “not enriched”). The statistical significance of the difference in clinical outcome was calculated using the log-rank Mantel-Cox test. The survival characteristics of the groups were visualized using Kaplan-Meier curves. *Corrplot* package (v0.84) was used to visualize the hazard ratio of each gene generated by the coxph() function of the survival package.

### Statistical analysis

Statistical comparisons were performed using *GraphPad Prism* v9.3.0 (RRID:SCR_002798; GraphPad Software, La Jolla, CA) or R. Values and error bars represent the mean ± standard error of the mean (SEM). Respective number of replicates (n) values are indicated in the figures or figure legends. *p*-values were determined by an appropriate statistical test such as Student’s t-tests or analysis of variance (ANOVA) with multiple comparison correction, as indicated in the figure legends. Power analyses were used to determine appropriate sample sizes for animal experiments (power 0.8, alpha 0.05).

### Data Availability

Sequencing data (bulk RNAseq, scRNAseq and ChIPseq) from this study will be publicly available without restrictions upon publication. The publicly available human gene expression profiling data sets of Cavalli *et. al.* (#GSE85217) (www.ncbi.nlm.nih.gov/geo/query/acc.cgi?acc=GSE85217), GSE28245 (www.ncbi.nlm.nih.gov/geo/query/acc.cgi?acc=GSE28245), and human Medulloblastoma microarray data from. St. Jude’s PeCan Data Portal (https://pecan.stjude.cloud/proteinpaint/yap1) were used to analyze expression levels of genes of interest in different MB subtypes. Publicly available YAP1 ChIPseq data from mouse heart tissue ^65^ (www.ncbi.nlm.nih.gov/geo/query/acc.cgi?acc=GSE115294, http://chip-atlas.org/view?id=SRX4159427, http://chip-atlas.org/view?id=SRX4159426) were used to analyze motifs associated with YAP1 binding sites.

### Code availability

All code to reproduce the figures presented in this manuscript will be available upon request from the authors.

## Acknowledgments

We thank Bikesh Nirala, Kin-Hoe Chow, Stephanie Wood, Hung Phan, Jia-Shiun Leu, and Active Motif for their technical assistance.

## Author contributions

KY: conceived the study, designed experiments, analyzed the data, and prepared manuscript

NA: designed experiments, and collected and analyzed the data and prepared manuscript SN: designed experiments, and collected and analyzed the data

JG: analyzed the sequencing data

JM, HNT, HB, RM: collected and analyzed data SC, FC, JO: provided reagents

All authors contributed to preparing the manuscript.

## Competing Interests

KY is a co-founder of EMPIRI, Inc. All other authors declare no competing interests.

**Supplementary Information is available for this paper.**

