## Supplementary Figures for "Sex-biased *Yap1* oncogene function"

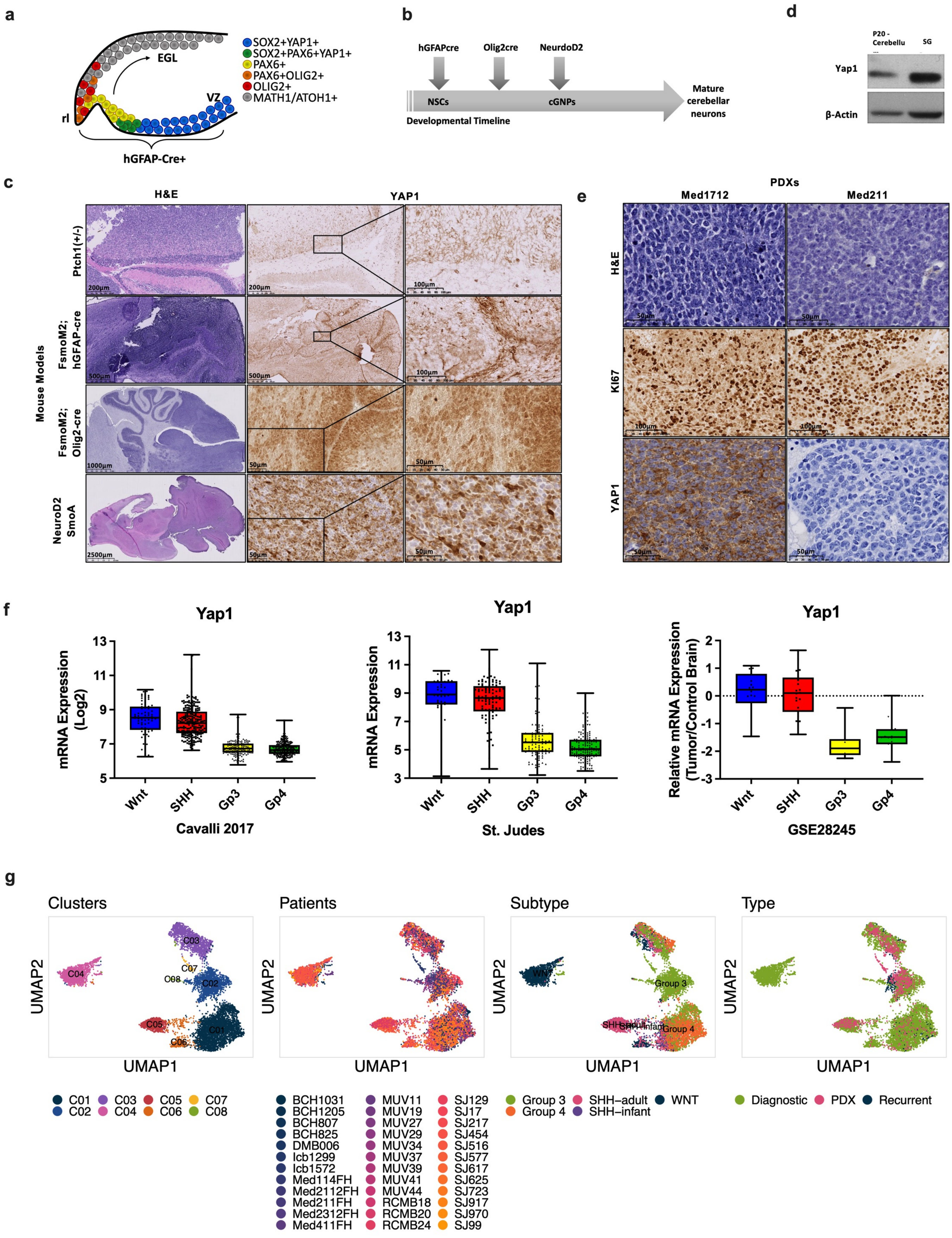
a
EGL
VZ
rl
hGFAP-Cre+
SOX2+YAP1+
SOX2+PAX6+YAP1+
PAX6+
PAX6+OLIG2+
OLIG2+
MATH1/ATOH1+
b
hGFAPcre
Olig2cre
NeurdoD2
NSCs
cGNPs
Developmental Timeline
Mature cerebellar neurons
d
P20 - Cerebellu
SG
Yap1
β-Actin
c
H&E
YAP1
Mouse Models
Ptch1(+/-)
FsmoM2; hGFAP-cre
FsmoM2; Olig2-cre
NeuroD2 SmoA
e
PDXs
Med1712
Med211
H&E
KI67
YAP1
f
Yap1
mRNA Expression (Log2)
Wnt
SHH
Gp3
Gp4
Cavalli 2017
mRNA Expression
St. Judes
Relative mRNA Expression (Tumor/Control Brain)
GSE28245
g
Clusters
Patients
Subtype
Type
UMAP1
UMAP2
C01 C02 C03 C04 C05 C06 C07 C08
BCH1031 BCH1205 BCH807 BCH825 DMB006 Icb1299 Icb1572 Med114FH Med2112FH Med211FH Med2312FH Med411FH
MUV11 MUV19 MUV27 MUV29 MUV34 MUV37 MUV39 MUV41 MUV44 RCMB18 RCMB20 RCMB24
SJ129 SJ17 SJ217 SJ454 SJ516 SJ577 SJ617 SJ625 SJ723 SJ917 SJ970 SJ99
Group 3 Group 4 SHH–adult SHH–infant WNT
Diagnostic PDX Recurrent

h

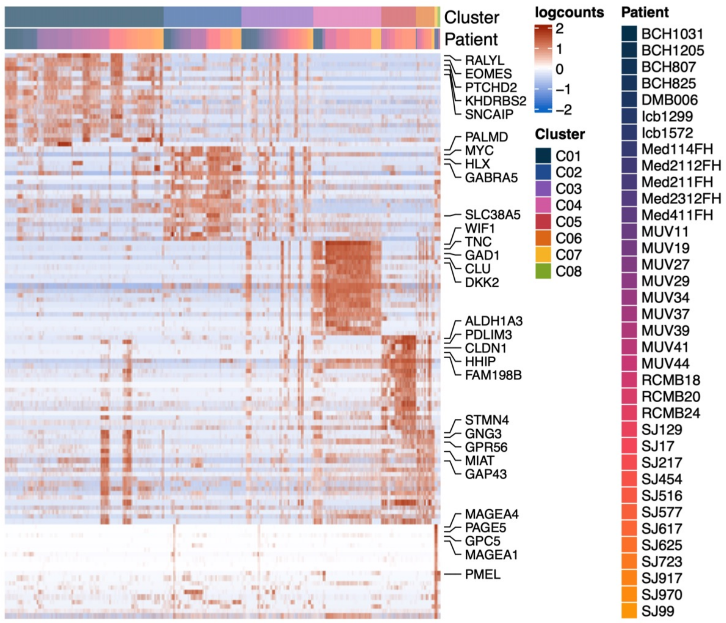


i

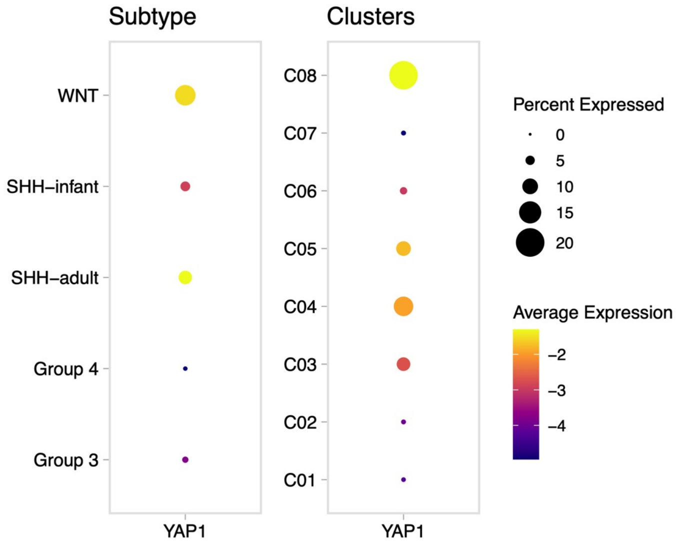


**Supplementary Figure 1: *Yap1* is reactivated in SHH MB regardless of cell of origin. a.** Schematic showing the different stages of differentiation of h*GFAPcre*-expressing cells. **b.** Schematic showing the different MB models on the developmental timeline. **c.** H&E stained section from *Ptch*$^{+/-}$, *fSMO-M2;hGFAPcre*, *fSMO-M2;Olig2Cre*, and *NeuroD2-SmoA* MB models in which the SHH pathway is activated in cerebellar cells at different stages of maturation. All models show YAP1+ cells. **d.** Western blot analysis showing overexpression of YAP1 in SG tumors compared with p20 cerebellum. **e.** IHC IHC sections from MB PDXs (MED1712 (SHH MB) and MED211 (group 3 MB) showing Ki67 and YAP1 staining. **f.** Meta-analysis of *YAP1* expression in human MB datasets (Cavalli et al. GSE85217), St. Jude's PeCan Data Portal, and GSE28245) showing higher expression of *Yap1* in WNT and SHH MB subtypes. **g.** Meta-analysis of external human MB dataset from Hovestadt *et al.* UMAP projections of 8691 cells aggregated from 36 samples (GSE119926), color-coded by identified clusters, sample ID, molecular subtype, and sample type. **h.** Top 20 differentially-expressed genes between clusters, ranked by FDR, are shown in the heatmap. Gene expression values were centered, scaled, and transformed to a scale from -2 to 2. Select signature genes are highlighted on the right and broad cell type assignment labels are on the right. **i.** Dot plot showing the expression of *YAP1*. Dot sizes indicate the percentage of cells in each cluster expressing the gene and colors indicate average expression.

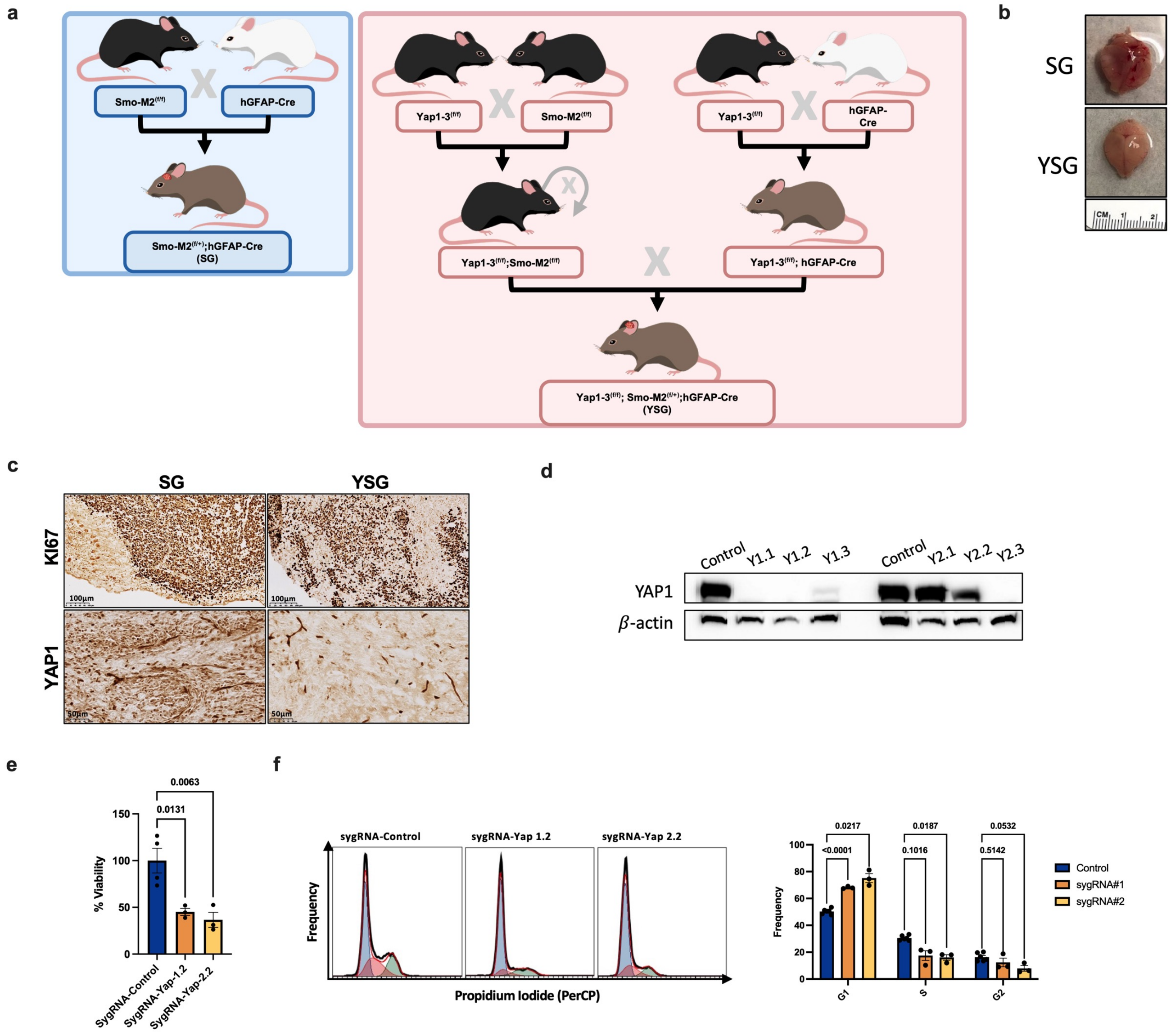


**Supplementary Figure 2: *Yap1* is activated in SHH medulloblastomas. a.** Schematic showing the breeding scheme for the generation of SG and YSG mice. **b.** Photograph of SG (p31-male) and YSG (p35-male) mice and their brains. **c.** IHC-stained sections from p20 SG and YSG tumors showing Ki67 and YAP1 staining. All experiments were performed with at least n=3. *P*-values were calculated using two-tailed Student's *t*-test. **d.** WB analysis of YAP1 expression in the mouse tumors. **e**. Viability of *Ptch;p53* CRISPR-Cas9 clones, control sygRNA vs two different sygRNAs targeting Yap1. n=3 each. *P*-values were calculated using one-way ANOVA followed by Dunnett's multiple comparisons test. **f**. Cell cycle analysis of *Ptch;p53* CRISPR-Cas9 clones, control sygRNA vs two different sygRNAs targeting Yap1. n=3 each. *P*-values were calculated using two-way ANOVA followed by Dunnett's multiple comparisons test.

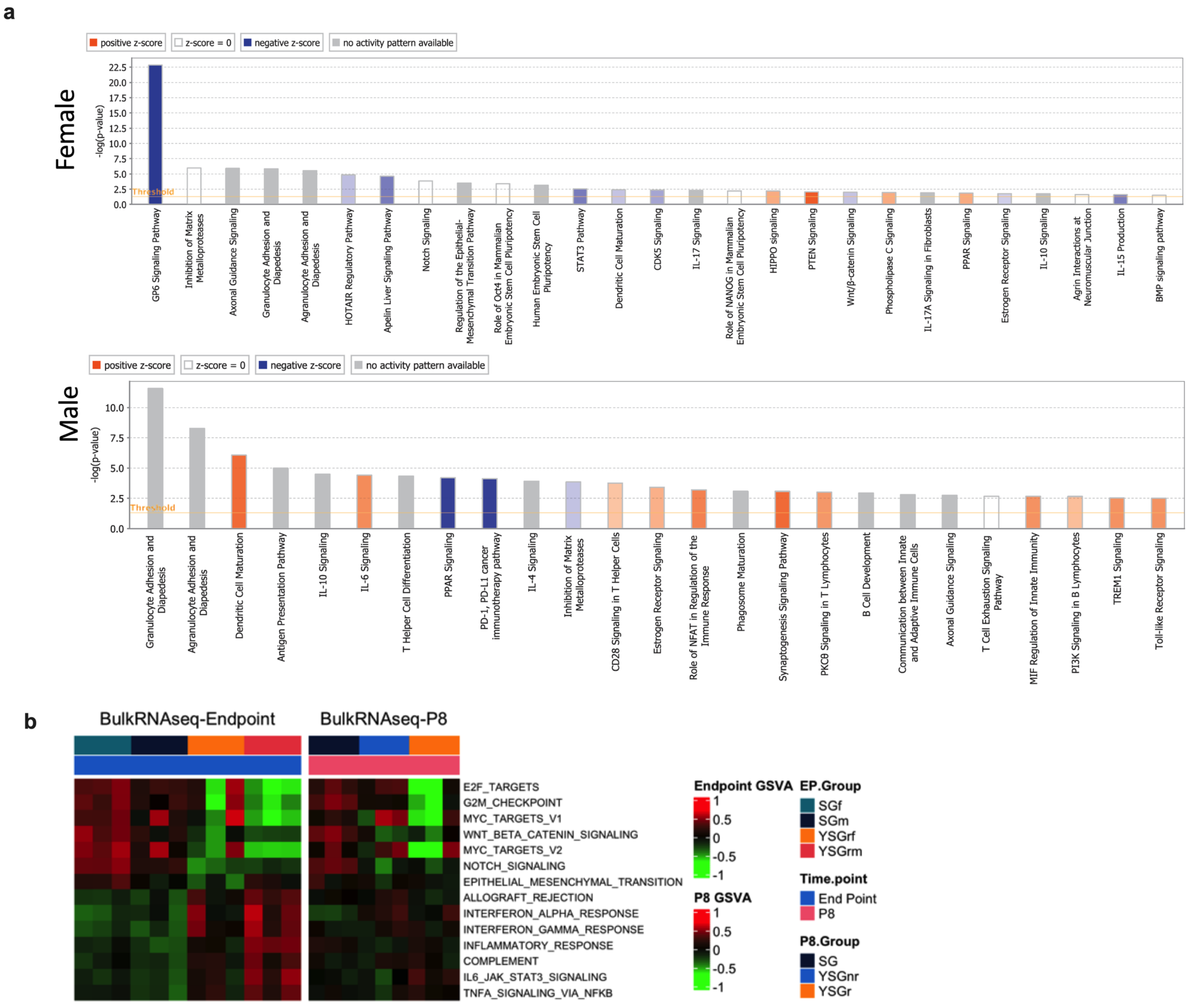


**Supplementary Figure 3: Bulk RNA-seq pathway analysis. a.** IPA analysis of differentially enriched canonical pathways across YSG$^r$ vs SG female (top) or male (bottom) tumor clusters. **b.** Gene Set Visualization Analysis (GSVA) of differentially enriched Hallmark pathways between endpoint YSG$^r$ and SG tumors (FDR <0.05). Same pathways are shown in bulk RNA-seq from p8 cerebella (right).

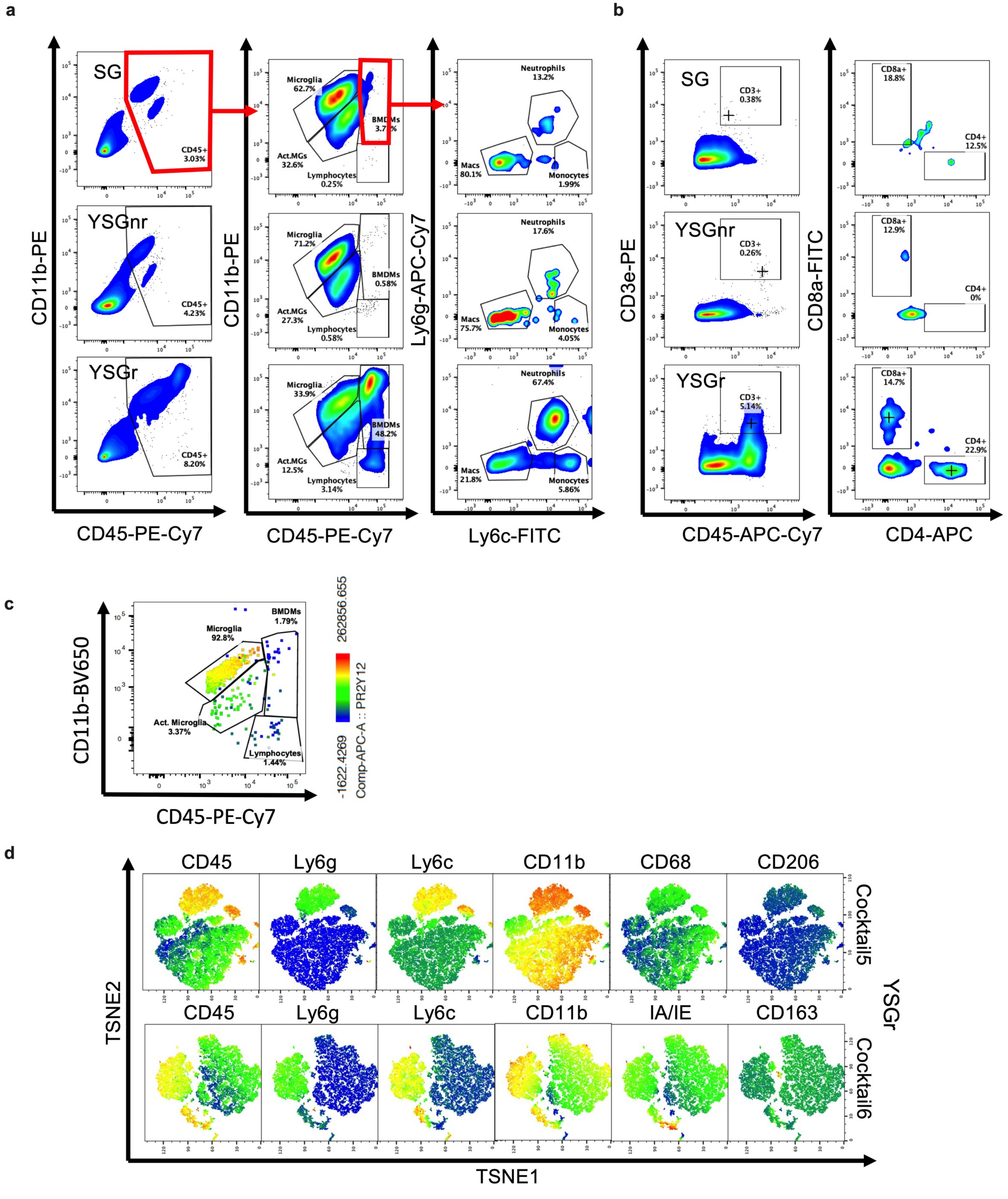


**Supplementary Figure 4: *Yap1* deletion increases bone marrow-derived monocyte immune infiltration in rescued SG medulloblastoma. a-b.** Representative flow cytometric analysis showing tumor-infiltrating BMDMs (**a**) or T cells (**b**) in SG, YSG$^{r}$, and YSG$^{nr}$ mice. Figures represent age- and sex-matched samples. **c.** Representative flow cytometric analysis showing P2RY12 expression in CD45 intermediate microglia. **d.** TSNE plots showing representative flow cytometric analysis of CD45$^{+}$ cells in a female YSG$^{r}$ tumors at p30. Color heatmap represents overlaid marker staining intensity (red=high, green=intermediate, blue=low).

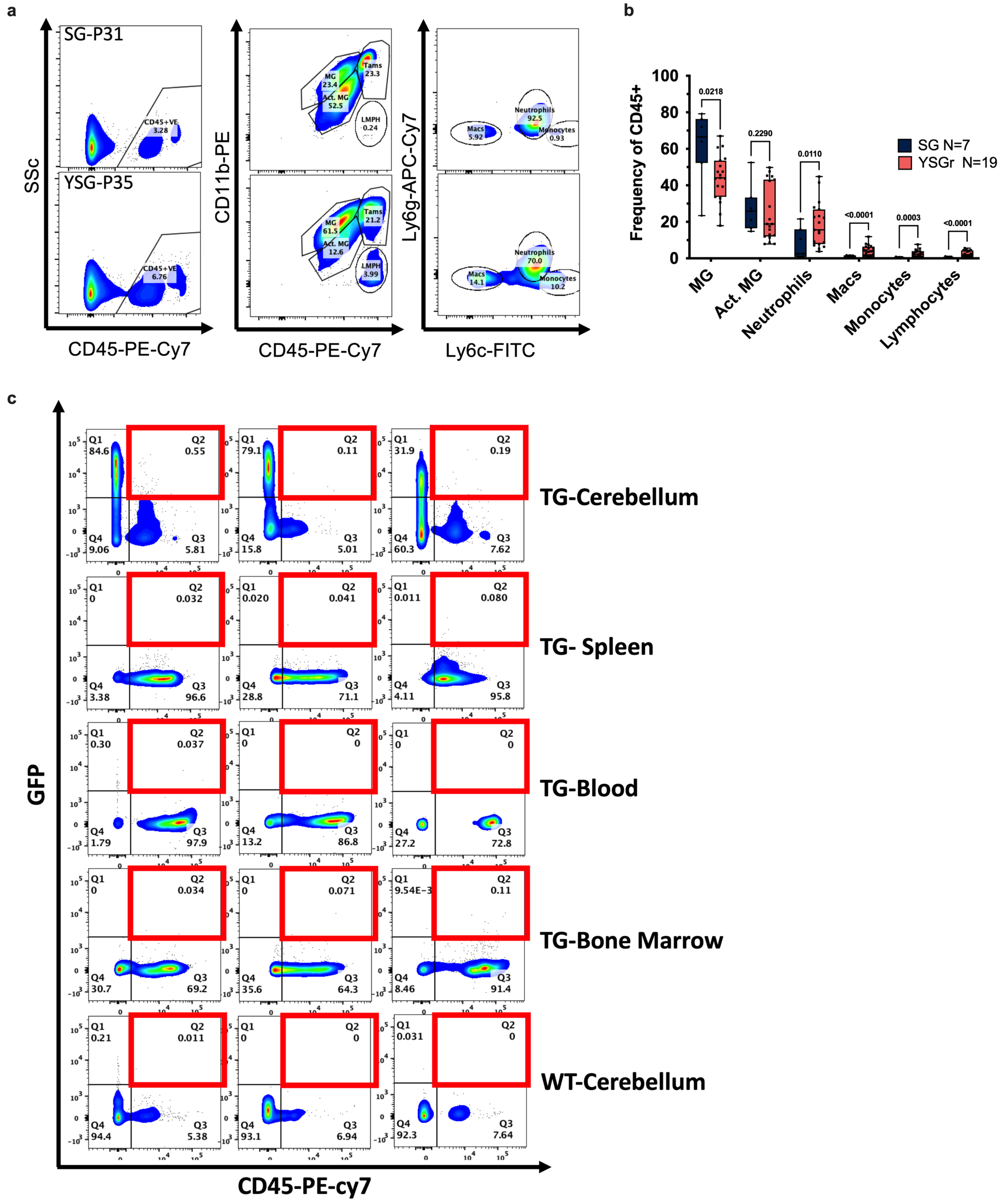


**Supplementary Figure 5: Effect of age on MB Immune infiltration. a.** Representative flow cytometric analysis of age-matched male SG (p31) and male YSG$^r$ (p35) samples. **b.** Quantification of flow cytometry analysis of overall post-weaning SG vs YSG$^r$. Number of mice is indicated in graph legend. Line inside box indicates median values. *P*-values were calculated using two-way ANOVA followed by the two-stage linear step-up procedure of Benjamini, Krieger, and Yekutieli. **c.** Flow cytometric analysis showing expression of GFP$^+$ cells in *ROSA26-fEGFP;hGFAPcre* cerebellum, spleen, blood and bone marrow.

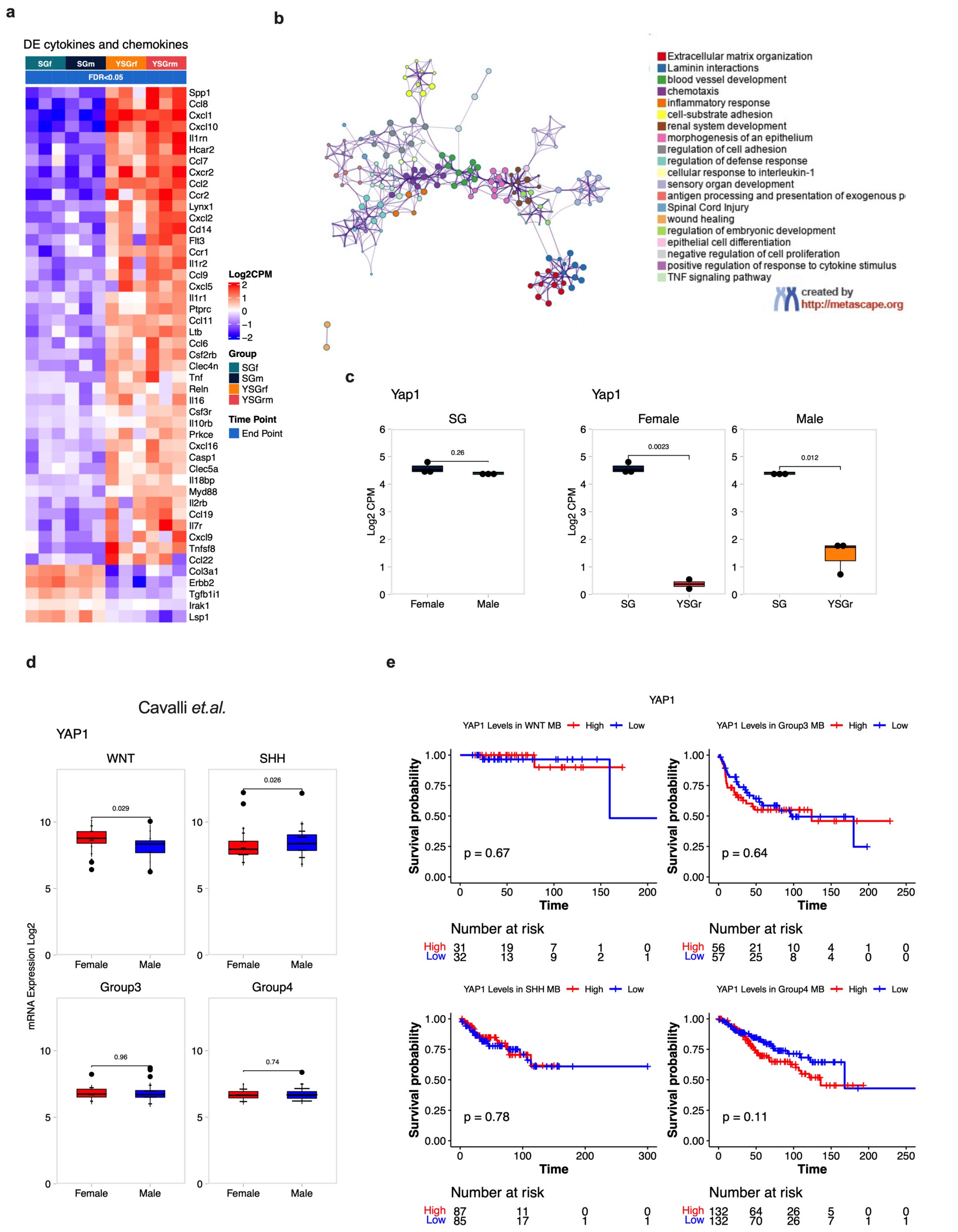


**Supplementary Figure 6: Additional bulk RNA-seq analysis. a.** heatmap showing row-centered log-CPM of differentially expressed cytokines, chemokines, and immune regulators in YSG$^r$ vs SG samples (FDR <0.05). **b.** Metascape analysis of DE genes between YSG and SG male and female samples showing Gene Ontology (GO) network. Each term is represented by a circle node, where its size is proportional to the number of input genes falling into that term, and its color represents its cluster identity. Terms with a similarity score >0.3 are linked by an edge (the thickness of the edge represents the similarity score). One term from each cluster is selected to have its term description shown as label. **c.** Box plots showing expression of *Yap1* in SG and YSG$^r$ bulk endpoint RNA-seq. **d.** Box plots showing expression of *YAP1* in human MB Cavalli *et al.* (GSE85217) dataset. **e.** Kaplan-Meier survival curve analyses of human MB Cavalli *et al.* (GSE85217) samples by subtypes. *YAP1* high and low expression cutoff was set at the median. *P*-values were calculated using the log-rank (Mantel-Cox) test.

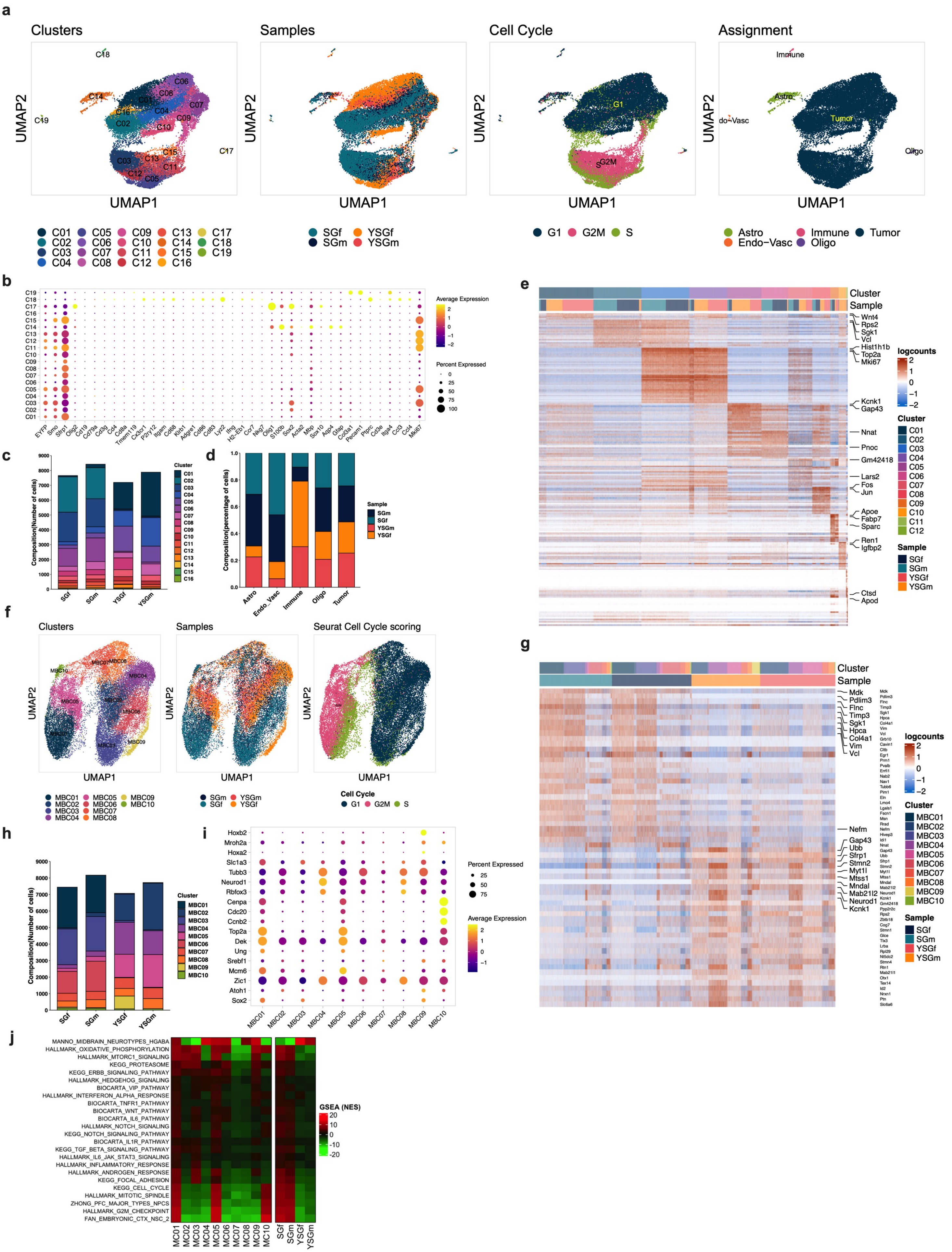

a
Clusters
Samples
Cell Cycle
Assignment
UMAP1
UMAP2
C01 C02 C03 C04 C05 C06 C07 C08 C09 C10 C11 C12 C13 C14 C15 C16 C17 C18 C19
SGf SGm YSGf YSGm
G1 G2M S
Astro Endo−Vasc Immune Oligo Tumor
b
Average Expression
Percent Expressed
EYFP Smo Sfrp1 Olig2 Cd19 Cd79a Cd3g Cd4 Cd8a Tmem119 Cx3cr1 P2ry12 Itgam Cd68 Klrb1 Adgre1 Cd86 Cd83 Lyz2 Ifng H2−Eb1 Ccr7 Nkg7 Olig1 S100b Sox2 Acta2 Mbp Sox10 Aqp4 Gfap Col3a1 Pecam1 Ptprc Cd3e Itga4 Cd3 Cd4 Mki67
c
Composition(Number of cells)
Cluster
d
Composition(percentage of cells)
Sample
Astro Endo_Vasc Immune Oligo Tumor
e
Cluster
Sample
logcounts
Wnt4 Rps2 Sgk1 Vcl Hist1h1b Top2a Mki67 Kcnk1 Gap43 Nnat Pnoc Gm42418 Lars2 Fos Jun Apoe Fabp7 Sparc Ren1 Igfbp2 Ctsd Apod
f
Clusters
Samples
Seurat Cell Cycle scoring
MBC01 MBC02 MBC03 MBC04 MBC05 MBC06 MBC07 MBC08 MBC09 MBC10
Cell Cycle
g
Cluster
Sample
logcounts
Mdk Pdlim3 Flnc Timp3 Sgk1 Hpca Col4a1 Vim Vcl Nefm Gap43 Ubb Sfrp1 Stmn2 Myt1l Mtss1 Mndal Mab21l2 Neurod1 Kcnk1
h
Composition(Number of cells)
Cluster
i
Hoxb2 Mrch2a Hoxa2 Slc1a3 Tubb3 Neurod1 Rbfox3 Cenpa Cdc20 Ccnb2 Top2a Dek Ung Srebf1 Mcm6 Zic1 Atoh1 Sox2
Percent Expressed
Average Expression
j
MANNO_MIDBRAIN_NEUROTYPES_HGABA
HALLMARK_OXIDATIVE_PHOSPHORYLATION
HALLMARK_MTORC1_SIGNALING
KEGG_PROTEASOME
KEGG_ERBB_SIGNALING_PATHWAY
HALLMARK_HEDGEHOG_SIGNALING
BIOCARTA_VIP_PATHWAY
HALLMARK_INTERFERON_ALPHA_RESPONSE
BIOCARTA_TNFR1_PATHWAY
BIOCARTA_WNT_PATHWAY
BIOCARTA_IL6_PATHWAY
HALLMARK_NOTCH_SIGNALING
KEGG_NOTCH_SIGNALING_PATHWAY
BIOCARTA_IL1R_PATHWAY
KEGG_TGF_BETA_SIGNALING_PATHWAY
HALLMARK_IL6_JAK_STAT3_SIGNALING
HALLMARK_INFLAMMATORY_RESPONSE
HALLMARK_ANDROGEN_RESPONSE
KEGG_FOCAL_ADHESION
KEGG_CELL_CYCLE
HALLMARK_MITOTIC_SPINDLE
ZHONG_PFC_MAJOR_TYPES_NPCS
HALLMARK_G2M_CHECKPOINT
FAN_EMBRYONIC_CTX_NSC_2
GSEA (NES)

**Supplementary Figure 7: scRNA-seq analysis of SG and YSG$^r$ tumors. a.** UMAP projections of 30,928 aggregate single cells from SG male, SG female, YSG$^r$ male, and YSG$^r$ female samples showing the composition of different cell types in mouse SHH MB. UMAP projections representing color-coded clusters by cluster numbers, cell origin by sample, cell cycle status, and assigned broad cell types. **b.** Dot plot showing the expression of marker genes for different cell types (medulloblastoma, astrocytes, endothelial cells, oligodendrocytes, and immune cells). Dot sizes indicate the percentage of cells in each cluster expressing the gene, and colors indicate average expression. **c.** Bar plots showing number of cells per sample colored by cluster. **d.** Bar plots showing percentage of cells in each assignment colored by sample. **e.** Top 20 differentially expressed genes in 12 clusters, ranked by FDR, are shown in the heatmap. Gene expression values were centered, scaled, and transformed to a scale from -2 to 2. Select signature genes are highlighted on the right. **f.** UMAP projections of 30,010 MB tumor cells only, color-coded by identified clusters, sample ID, and cell cycle status. MB cells in clusters 1:8 from Figure 1a were extracted and analyzed through *de novo* clustering. **g.** Top 30 differentially expressed genes between samples in tumor cells only, ranked by FDR, are shown in the heatmap. Gene expression values were centered, scaled, and transformed to a scale from -2 to 2. Select signature genes are highlighted on the right, and broad cell type assignment labels are on the right. **h.** Bar plots showing number of tumor cells per sample colored by cluster. **i.** Dot plot showing the expression of neuronal markers. Dot sizes indicate the percentage of cells in each cluster expressing the gene, and colors indicate average expression.

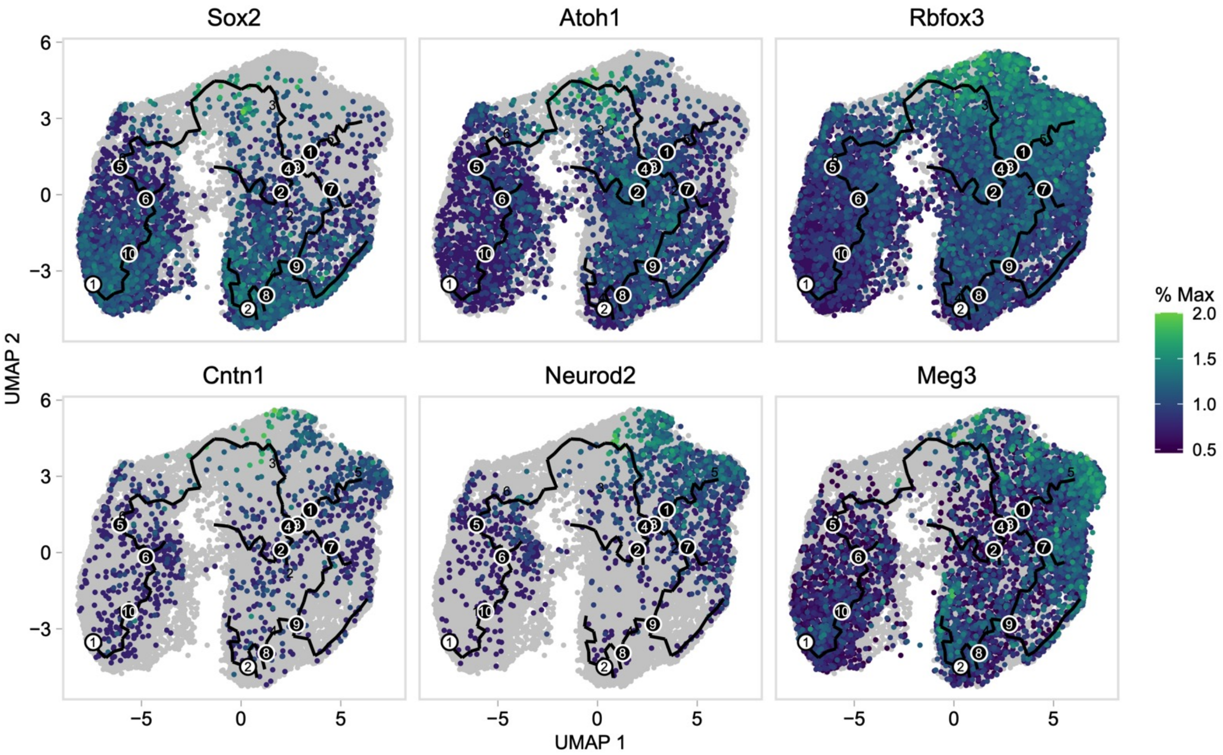


**Supplementary Figure 8: YSG$^{r}$ tumors show increased signs of differentiation.** Feature plots showing expression of major lineage markers.

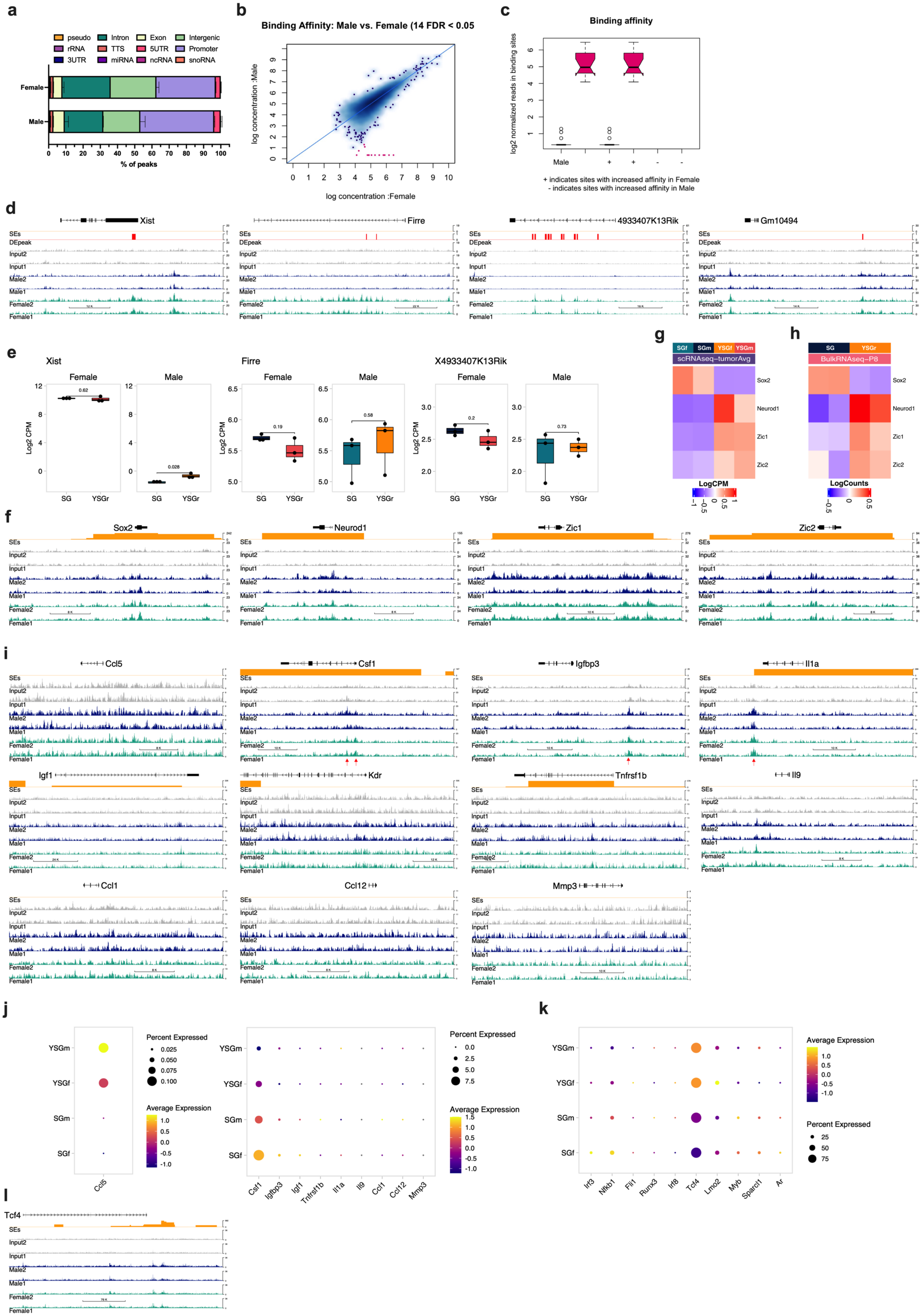
a
pseudo
Intron
Exon
Intergenic
rRNA
TTS
5UTR
Promoter
3UTR
miRNA
ncRNA
snoRNA
Female
Male
% of peaks
b
Binding Affinity: Male vs. Female (14 FDR < 0.05
log concentration :Male
log concentration :Female
c
Binding affinity
log2 normalized reads in binding sites
Male
+ indicates sites with increased affinity in Female
- indicates sites with increased affinity in Male
d
Xist
Firre
4933407K13Rik
Gm10494
SEs
DEpeak
Input2
Input1
Male2
Male1
Female2
Female1
e
Xist
Firre
X4933407K13Rik
Female
Male
Log2 CPM
SG
YSGr
g
SGf
SGm
YSGf
YSGm
scRNAseq–tumorAvg
Sox2
Neurod1
Zic1
Zic2
LogCPM
h
SG
YSGr
BulkRNAseq–P8
LogCounts
f
Sox2
Neurod1
Zic1
Zic2
i
Ccl5
Csf1
Igfbp3
Il1a
Igf1
Kdr
Tnfrsf1b
Il9
Ccl1
Ccl12
Mmp3
j
YSGm
YSGf
SGm
SGf
Ccl5
Percent Expressed
Average Expression
Csf1
Igfbp3
Igf1
Tnfrsf1b
Il1a
Il9
Ccl1
Ccl12
Mmp3
k
Irf3
Nfkb1
Fli1
Runx3
Irf8
Tcf4
Lmo2
Myb
Sparcl1
Ar
l
Tcf4

**Supplementary Figure 9: Direct targets of YAP1 in MB. a.** The location of YAP1 peaks relative to genomic annotations is presented in a stacked bar plot. Error bars represent SEM. **b.** MA plot showing differential binding affinity between female and male samples. Each dot represents a binding site. 14 red dots represent differentially bound sites identified by edgeR (FDR <0.05, fold difference >2). **c.** Boxplots showing read distribution amongst all differentially bound sites (left, n=14), sites in female samples (middle, n=14), sites in male samples (right, n=0). **d.** YAP1 binding sites to genes identified by edgeR as differentially bound in female vs male samples. **e.** Boxplots showing the expression of *Xist*, *Firre*, and *4933407K13Rik* in endpoint bulk RNA-seq samples.**f.** YAP1 binding sites in *Sox2, Neurod1, Zic1*, and *Zic2* genomic regions. **g.** Heatmap showing expression of *Sox2, Neurod1, Zic1*, and *Zic2* in scRNA-seq **h.** Heatmap showing expression of *Sox2, Neurod1, Zic1*, and *Zic2* in P8 bulk RNAseq datasets. **i.** YAP1 binding sites in RANTES/*Ccl5,* M-CSF*/Csf1*, *Igfbp3, Il1a, Igf1*, VEGF R2/*Kdr*, sTNF RII/*Tnfrsf1b*, *Il9*, TCA-3/*Ccl1,* MCP-5*/Ccl12*, and *Mmp3* genomic regions. **j.** Dot plot showing the expression of **Figure 3h** proteins. Dot sizes indicate the percentage of cells in each cluster expressing the gene and colors indicate average expression. **k.** Dot plot showing the expression of transcription factors that bind to *Ccl5* genomic regions. Dot sizes indicate the percentage of cells in each cluster expressing the gene, and colors indicate average expression. **l.** YAP1 binding site in *Tcf4* genomic region.

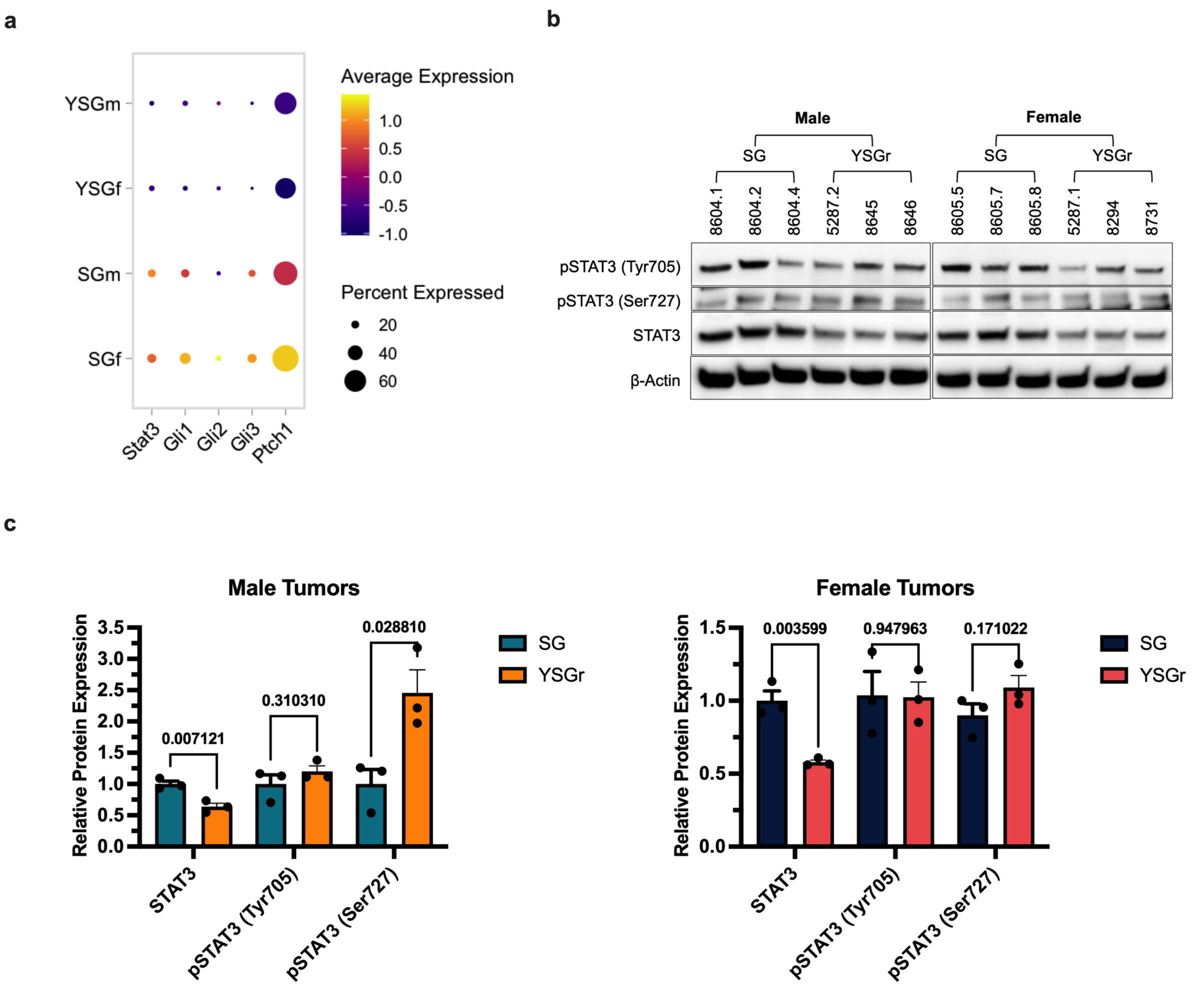


**Supplementary Figure 10: STAT3 expression is reduced in both male and female scRNA-seq MB tumor cells. a.** Dot plot showing the expression of *Stat3, Gli1, Gli2, Gli3*, and *Ptch1* in MB tumor cells only. Dot sizes indicate the percentage of cells in each sample expressing the gene, and colors indicate average expression. **b.** Western blot analysis of STAT3, pSTAT3(Ser727), and pSTAT3(Tyr705) protein levels in SG or YSG$^r$ male or female tumors. **c.** Quantification of western blot results from (b). All values were normalized to Actin control and then pSTAT3 values were additionally normalized to STAT3 values. *P*-values were calculated using Student's *t*-test.
